# A general model for the evolution of thermal performance curves with application to real time-series data

**DOI:** 10.64898/2026.06.17.733045

**Authors:** Jiseon Min, Zoe Chapman, Ellie McCabe, Joaquin C. B. Nunez, Nicholas Teets, Katie E. Lotterhos

## Abstract

Thermal performance curves (TPCs) are widely used to investigate the effect of changes in body temperature on an organism’s performance. Despite empirical evidence that temperature-dependent performance is ubiquitous across taxa, the field lacks models for how thermal performance evolves under realistic timeseries, genetic architectures, and physiological constraints. We address this gap by integrating a mathematical model with individual-based quantitative population genetic simulations. Our model can predict the evolutionary trajectory and shape of TPCs for any given thermal regime. Our model reproduces core properties of TPC evolution from previous studies such as the emergence of generalists in variable environments, but also explains why organisms may evolve TPCs that do not match their historical body temperature range. We uncover novel dynamics of adaptive tracking, the most notable being multi-generation lags between temperature and TPC parameters that can lead to unexpected correlations between the two. Our model predicts empirically observed patterns of adaptive tracking of critical thermal minimum in the invasive pest *Drosophila suzukii*, including individual-level variability and multi-generation lags with changing temperature. Our results also highlight the limitations of models that ignore factors that influence TPC evolution and individual variability, such as autocorrelation in temperature timeseries, effective population size, evolution of additive genetic correlation in TPC parameters, genetic architecture, and physiological constraints. Our flexible simulation model can incorporate these factors and help generate empirically testable hypotheses of how species will evolve in response to global climate change.

## Introduction

A central question in thermal biology is how to predict the performance of different species in response to changing temperatures from seasonal fluctuations, extreme events, and anthropogenic climate change [16]. Thermal performance curves (TPCs), a measure of physiological performance related to organisms’ fitness as a function of body temperature, are used by both empiricists and theorists to predict how species will respond to climate change [3, 24]. Various modeling approaches have been developed to forecast species responses to temperature change based on TPCs [11, 12, 20, 40]. However, these approaches do not account for the fact that there can be significant variation in thermal performance curves within a species, that variation may be heritable, and that TPCs can evolve as the population adapts to novel thermal environments (see ref [45] for a comprehensive review of common assumptions around TPCs). Overall, genetic adaptation is recognized as a crucial piece of information to predict species responses to warming, but currently no existing framework dynamically integrates environmental variability with corresponding evolution in the thermal performance curve. This leaves a gap in our ability to forecast how organisms will respond to increasingly variable thermal regimes under climate change.

Although some genetic models have been developed to study TPC evolution that treat the parameters of the TPC as heritable traits [15, 31, 42, 48], restrictive assumptions of these models limit our ability to use them for broader contexts. For instance, recent mathematical models such as [15] and [30] have assumed a non-evolvable genetic variance in TPC-related traits, which is an unrealistic assumption because genetic variance in TPC-related traits and the covariance among them evolve toward a new equilibrium over multiple generations due to finite population size, linkage disequilibrium, and a finite mutation rate [4]. Also, previous models have assumed unlimited mutational supply and have not imposed physiological limits on TPC evolution, which implies that TPCs can infinitely track climate change. However, empirical studies have highlighted the importance of physiological limits in predicting organism’s vulnerability to climate change [8, 21, 25].

The limitations of previous models also prohibit their use for studying seasonal adaptive tracking, which occurs when a population’s genetic changes continuously follow shifting thermal optima across seasons [31]. There exists empirical evidence for seasonal adaptive tracking of temperature with previous studies reporting rapid phenotypic and genotypic adaptation to temperature [6, 7, 33, 36, 41]; however, we lack models to explain the conditions under which polygenic thermal performance traits adaptively track temperature. Although some recent models have studied allele frequency fluctuations in seasonally fluctuating environments [49], these models have assumed alleles have direct effects on fitness in two discrete environmental states and have focused on whether or not switching dominance is necessary for genetic polymorphisms to be maintained in fluctuating environments [13, 14]. Thus, there is a need for a new model of seasonal adaptive tracking with more realistic genetic architectures and a more concrete connection to thermal physiology and thermal traits.

In this paper, we develop a novel analytical model that extends previous models by incorporating physiological constraints, which allows for the calculation of the population-mean TPC for a constant population temperature with a normal distribution of individual body temperatures. We then overcome the limitations of the analytical model by extending our framework to individual-level forward-time population genomic simulations that use real time-series of body temperature within and across generations as input. Our simulation approach allows us to explore how TPC evolution proceeds under linkage, drift, polygenicity, and evolving genetic variance and covariance among TPC parameters. While the analytical and simulation approaches converge in terms of some key predictions, such as mismatch between body temperature and expected thermal optimum, some features like the influence of finite population size and seasonal adaptive tracking were only possible through simulation, highlighting the benefit of explicit and realistic population genomic simulation.

We use this model to study the dynamics of seasonal adaptive tracking, and to explore conditions under which a population’s evolved TPC may not align with the historical body temperatures of the population. Specifically, we ask: (i) under what parameter space do we find a mismatch between the distribution of body temperatures and the evolved TPCs?; (ii) can our model predict empirically observed patterns of adaptive tracking of a thermal performance trait in the wild?; and (iii) how do different aspects of realism (e.g. seasonally fluctuating generation time, genetic drift, and autocorrelation in body temperature) affect the evolution of TPCs?

## Results

We derive a novel formula for the thermal performance curve (*w_T PC_*, Figure 1B) by combining the TPC formula by Deutsch et al. [19] (Figure 1A) with physiological constraints for the expression of extreme trait values for the critical thermal minimum (*CTmin*), breadth (*B*), and critical thermal maximum (*CTmax*)(Figure 1B; *w_CT min_, w_CT max_, w_B_*. See Figures S1 and S2 for more visual representations of each fitness). We use this formula to analytically model the fitness landscape (Figure 1D) for a given distribution of body temperature within an individual’s lifetime (Figure 1C). We analytically approximate the evolutionary trajectory to the evolved TPC using the selection gradient on the fitness landscape (Figure 1E, Method 1). We then extend this framework to novel simulations in a finite population in SLiM [22] for body-temperature timeseries with any given timestep, by treating the aforementioned TPC parameters as quantitative traits determined by the additive effects of individual mutations (Figure 1E, Method 2).

**Figure 1:**
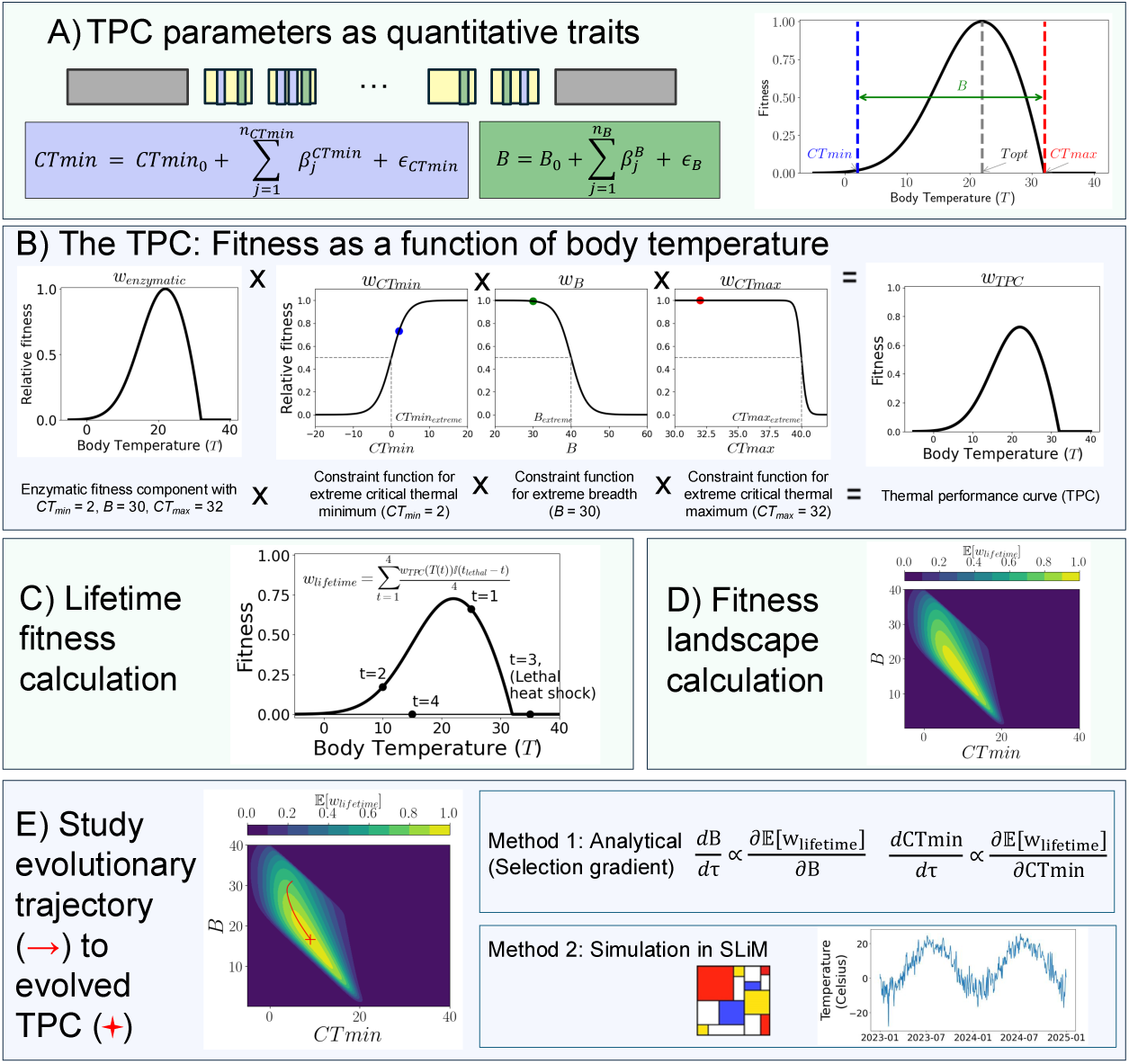
Overview of analytical and simulation approach. A) We model the parameters on the thermal performance curve (TPC): *B* is breadth, *CTmin* is critical thermal minimum, *Topt* is thermal optimum, and *CTmax* is critical thermal maximum. With the simplifying assumption that *Topt* = *CTmin* + 2*B/*3 and *CTmax* = *B* + *CTmin*, we reduce the theory to modeling two parameters as quantitative traits (*CTmin* and *B*). In simulations, each chromosome has neutral sites (grey; 20kb each) and 12 linkage blocks (yellow; 5kb each) where random QTN mutations can arise. Every new mutation affects either *B* (green) or *CTmin* (blue) and the effect size is drawn from a normal distribution with mean 0 and variance 0.05. For simplicity, we assume every mutation has effect on only one of the two key parameters. See Table S1 for information on evolved genomic architecture (number of segregating QTNs and distribution of effect sizes) in our simulations. Each diploid individual’s *CTmin* and *B* are determined from their default values (i.e. values of *CTmin* or *B* without QTN mutations or environmental noise; *CTmin*_0_*, B*_0_), sum of effect sizes of QTN mutations the individual carries (*β_CT min_, β_B_*), and environmental noise (*ɛ_CT min_, ɛ_B_*). B) The TPC is calculated from four relative fitness components: an enzymatic fitness function and three physiological constraint functions that prevent the system from evolving extreme TPC parameters. The function *w*_enzymatic_ describes fitness as a function of temperature (given *CTmin* and *B*), *w_CT min_* models declining fitness as *CTmin* becomes extremely low (*CTmin* ≪ *CTmin*_extreme_), *w_B_* models declining fitness as *B* becomes extremely large (*B* ≫ *B*_extreme_), and *w_CT max_* models declining fitness as *CTmax* becomes extremely high (*CTmax* ≫ *CTmax*_extreme_). We define critical physiological thresholds of each of the TPC parameters as the parameter value at which relative fitness is 0.5. Multiplying these fitness components together incorporates a physiological cost into the TPC for a given set of parameters and gives *w_T PC_*: the thermal performance curve. See Figures S1 and S2 for more visual representations of each fitness component. C) The lifetime fitness of an individual takes into account variability in body temperature at each timestep and thus the variability in fitness during its lifetime. The lifetime fitness *w*_lifetime_ is calculated as the average of fitness at each timestep *t*, given by *w_T PC_* for temperature *T* at the timestep or 0 after *t*_lethal_ (the first day when body temperature exceeds the lethal limit). An example with generation length equal to four timesteps is shown for an individual that is exposed to lethal heat-shock at *t* = 3; its fitness at subsequent timesteps is 0, but its lifetime fitness is non-zero and calculated as the mean fitness over the entire lifetime. In our SLiM simulations, which uses a Wright-Fisher model, *w*_lifetime_ is used as weights for binomial sampling of new set of individuals in the subsequent generation. D) The expected fitness *E*[*w*_lifetime_] for a given probability distribution of body temperature is calculated for all possible values of *B* and *CTmin* to create the expected fitness landscape. See SI-methods for derivation of *E*[*w*_lifetime_]. E) The evolutionary trajectory to the evolved TPC is then studied with an analytical approach or simulations. With the analytical approach, the trajectory of a trait over an arbitrary time *τ* is estimated as the partial derivative of the fitness landscape *E*[*w*_lifetime_] with respect to that trait. With SLiM simulations, the trajectory can be studied given any timeseries of body temperature, generation length, and other evolutionary parameters (genetic map, recombination rate, mutation rate, etc.).

### TPC evolution in constant environments

We first consider a simplified example where individuals reside in a constant environment; each individual samples its body temperature from a normal distribution with the population mean (5, 20, or 35 *^◦^*C) and standard deviation in individual body temperature (1 or 3 *^◦^*C) each day. We find that the analytical theory (see SI-Method) accurately predicts the average simulated trajectory of the population-mean TPC as long as selection dominates over drift (SI Results C; Figure S3 and S4).

For the simulations, we chose a relatively high mutation rate for quantitative genetic nucleotides (QTNs) that resulted in a highly polygenic architecture, with several hundreds of causal mutations segregating at the end of the simulation (Table S1). All simulations reached equilibrium by about 20k-30k generations (Table S2). Each panel in Figure 2(a) shows 40 randomly sampled individuals’ *w_T PC_* from a single simulation for a given distribution of body temperatures. The thermal optimum evolves with mean body temperature (*Topt*, left to right columns in Figure 2(a)), and that higher variance leads to broader TPCs (i.e. bigger *B*, top to bottom rows in Figure 2(a)). The panels in Figure 2(b) show the analytical fitness landscapes for two different distributions of body temperatures and the individual simulated TPCs. For example, more variable body temperatures evolve a lower *CTmin* and higher breadth, while tropical body temperatures evolve a higher *CTmin* and narrower breadth Figure 2(b).

**Figure 2:**
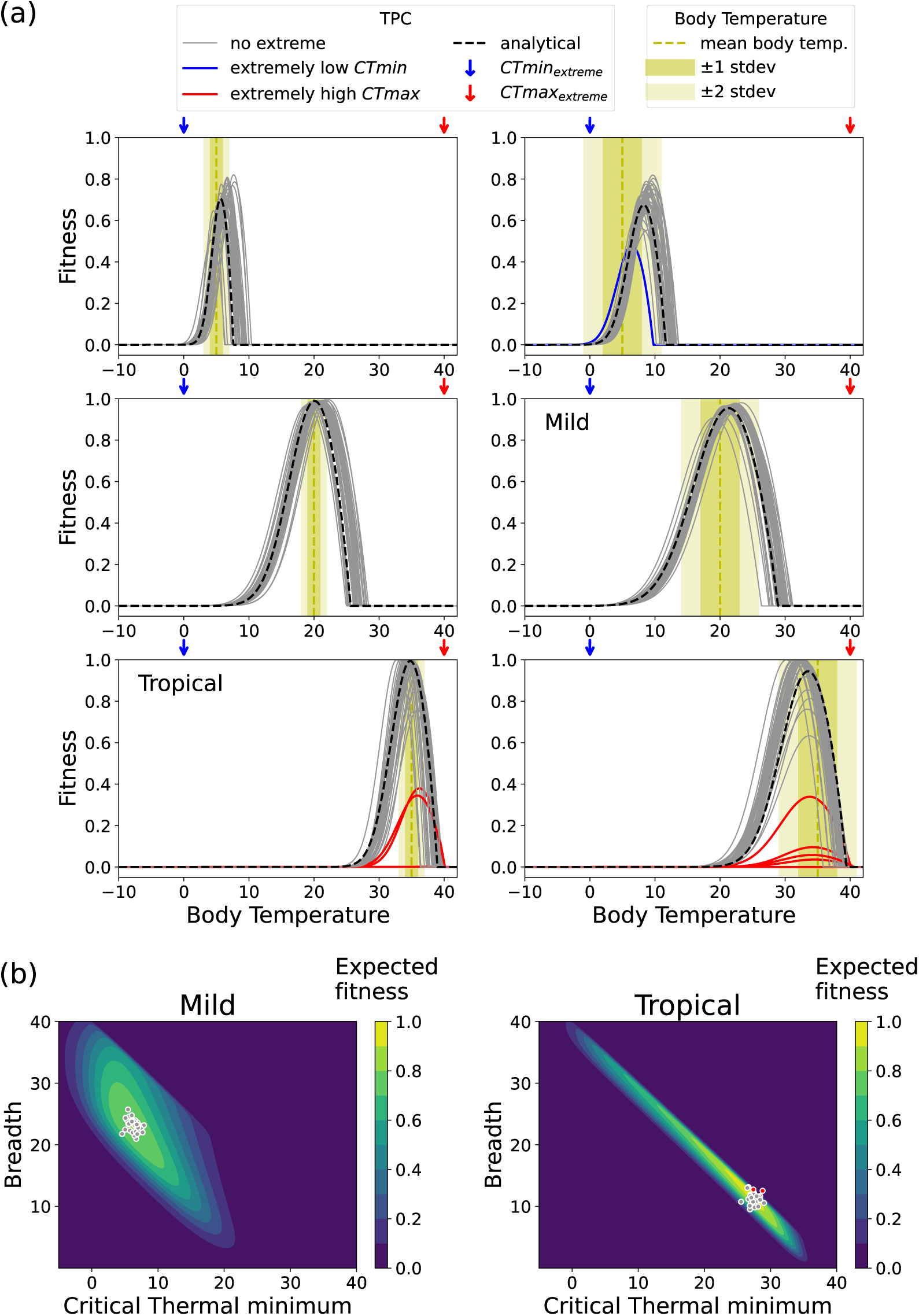
The evolution of thermal performance curves (TPCs) distribution of body temperatures (yellow background) around a constant population-mean temperature (vertical yellow dashed line). (a) Comparison of the analytical population-level TPC (black dash line) and simulated individual TPCs (grey, blue, and red lines). Note the mismatch in some panels between the body temperature distribution and the evolved TPCs. The color of the TPC indicates whether an individual has an extreme TPC parameter(s) that incurs a fitness cost: “no extreme” TPCs experience negligible fitness penalty from physiological constraints, “extremely low *CTmin*” TPCs have *CTmin* below or at the *CTmin*_extreme_ = 0*^◦^*C, and “extremely high *CTmax*” TPCs have *CTmax* above or at the *CTmax*_extreme_ = 40*^◦^*C. (b) Fitness landscapes for two different distribution of body temperatures (“mild” (left) and “tropical” (right) environment, as labeled in top panel). Individual simulated TPCs (circles, colored based on same criteria as top panel) are found near the fitness maximum on analytical fitness landscapes (*E*[*w*_lifetime_]). See Methods for parameters used in simulation.

Our simulations reveal interesting patterns of individual variability in TPCs that evolve within a population. Overall, we find good agreement between the analytical fitness landscape and the simulated TPCs, but individuals with extreme TPCs do evolve in some scenarios (SI Results B; Figure 2(a)). For instance, in the hottest and most variable temperature condition (lower right corner of Figure 2(a)), the population is regularly subjected to heat stress greater than the critical value of *CTmax* (greater than 40*^◦^*C as shown by the yellow body temperature distribution in the background). In this scenario, a small proportion of individuals evolve TPCs with extreme *CTmax* values beyond the critical threshold (red TPCs). Although individuals with these TPCs incur a physiological cost (as evidenced by the lower maximum fitness of their TPCs), these extreme traits persist at low frequency in the population for two reasons: because these individuals have the highest fitness at extreme body temperatures, and because of phenotypic noise that was modeled in the expression of the trait (i.e. *ɛ_CT min_* and *ɛ_B_* in Figure 1A).

#### Under what parameter space do we find a mismatch between the distribution of body temperatures and the evolved TPCs?

Our model also reveals interesting dynamics about the evolution of the thermal breadth and thermal optimum and its mismatch with distribution of body temperatures, with relevance to species responses to climate change. In conditions characterized by a narrow distribution of body temperatures that *do not* exceed physiological extremes, TPCs evolve to have wider breadth than expected (e.g. Figure 2(a), middle left panel, note wider TPCs than body temperature) because there are no costs. As the distribution of body temperatures becomes wider but still does *not* exceed *CTmax*_extreme_, the thermal optimum evolves to exceed the mean body temperature because of the steeper fitness decline on the right side of the TPC, consistent with previous studies [48] (e.g. Figure 2(a), top right and middle right panels, note higher *Topt* than mean body temperature). Finally, in conditions characterized by a wide distribution of body temperatures that *does* exceed *CTmax*_extreme_, TPCs evolve to have narrower breadths than expected and lower thermal optimum than the mean body temperature (e.g. Figure 2(a), lower right panel) owing to the fitness cost associated with expressing extreme *CTmax*. In the SI (SI Results A; Figure S5), we extend our analysis to bimodal distributions and find even greater mismatch between the body temperature distribution and thermal optimum as the population effectively forsakes the harder-to-adapt-to thermal regime.

#### On the evolved correlation among TPC parameters

Ultimately, the evolved correlation of TPC parameters will be determined by the balance between multiple forces of evolution including the fitness landscape (Figure 2b), selective pressures from hot and cold body temperatures, linkage disequilibrium, and genetic drift. Although we set the effects on *CTmin* and *B* from each new mutation to be independent in simulations (i.e. diagonal mutational variance-covariance matrix; see Methods), negative correlations evolve between *CTmin* and *B* (Figure S6 and bottom row of Figure S7) due to the negative correlation on the fitness landscape (Figure 2b). The degree of correlation between *CTmin* and *B* for a given fitness landscape, however, is more strongly impacted by genetic drift: smaller populations have weaker correlations (Figure S8). This result challenges previous simulation studies that assumed a fixed genetic correlation between TPC parameters and underscores the need for explicit individual-based simulations [15]. Overall, our simulation suggests that the strength of correlation between evolved TPC parameters can wildly differ based on the range of temperatures experienced by the population and population size (see SI results D).

### TPC evolution in seasonally fluctuating environments

Next, we let the mean body temperature change seasonally while not adding any special parameterization (e.g. switching dominance as in ref [49]) and investigate whether seasonal adaptive tracking emerges solely from mean body temperature oscillating between generations.

#### Adaptive tracking to sinusoidal temperature

We first simulate a population experiencing a sinusoidal fluctuation in body temperature between 0 and 35*^◦^*C with a small Gaussian noise added for each individual every day (stdev = 1*^◦^*C) and with a 360 day period (i.e. 1 year = 360 days; Figure 3(a), gray dashed line). For simplicity, each generation is 10 days long (see Methods). Each simulation lasted for 20k generations (roughly 550 years) including a 5k-generation burn-in period.

**Figure 3:**
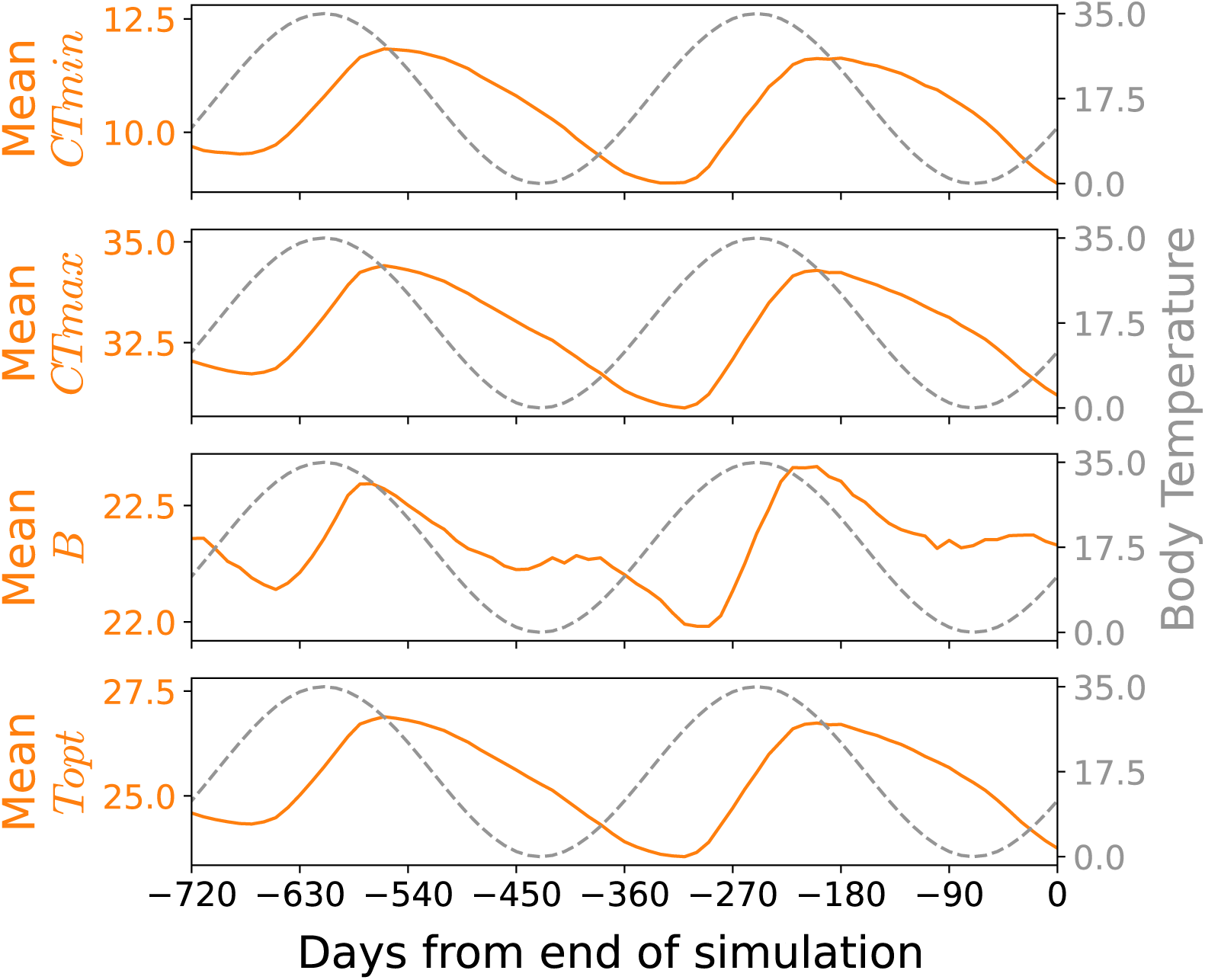
TPC parameters oscillate in response to sinusoidally fluctuating mean body temperature. All parameters of *w_T PC_* are set to default values (see Methods) except for *B*_extreme_ which is lowered to 20 for better visualization – see Figure S11 for results from changing the hyperparameters. As daily body temperature fluctuates sinusoidally between 0 and 35 with period of 360 days with small individual fluctuations (variance = 1), all four key TPC parameters adaptively track the change with a lag. The lag is shorter following summer than winter.

All TPC parameters in these simulations adaptively tracked body temperature, but with varying temporal lags (see SI Methods for how lag times were measured). *CTmin* oscillated with a shorter lag in summer and a longer lag in winter: it is minimized about 110 days (= 11 generations) after peak winter and maximized after approximately 63 days (=6.3 generations) after the peak summer (Figure 3 top).

In contrast, *CTmax* had slightly different lags than *CTmin*: it reached its minimum after about 120 days (=12 generations) after peak winter and its maximum about 51 days (=5.1 generations) past peak summer (Figure 3, second row). The different lags in *CTmax* and *CTmin* causes a distinct bimodal waveform in *B* (= *CTmax* − *CTmin*) where it rapidly approaches a yearly maximum shortly after peak summer, briefly increases again shortly after peak winter, and then is minimized in the spring (Figure 3, third row). The faster adaptation in *CTmax* following summer than in *CTmin* winter is consistent with higher genetic variance in fitness in summer, which leads to more rapid adaptation according to Fisher’s fundamental theorem (SI Results E; Figure S9 and S10).

#### Can our model predict empirically observed patterns of adaptive tracking of a thermal performance trait in the wild?

*D. suzukii*, an invasive fruit fly species native to South Asia, has spread rapidly in North America with more extreme winters, raising questions about the ability of their cold tolerance to genetically adapt. We compared our model predictions to the patterns observed in a recent study that sampled wild populations of *D. suzukii* in the summer and fall over multiple years and discovered seasonal adaptive tracking in *CTmin* [34] (Figure 4A). One counterintuitive feature of these data is that flies sampled at the peak of summer exhibit the lowest *CTmin*, yielding a negative correlation with body temperature at zero time lag, with the correlation becoming positive at a lag of 60 to 100 days (4-8 generations) (Figure 4C). This pattern has been proposed to be a legacy of overwintering: flies collected early in the season are descended from winter survivors and initially retain high cold tolerance, which then erodes over the summer [44]. However, despite this biological explanation, the dynamics governing the timing and magnitude of these lagged responses remain understudied.

**Figure 4:**
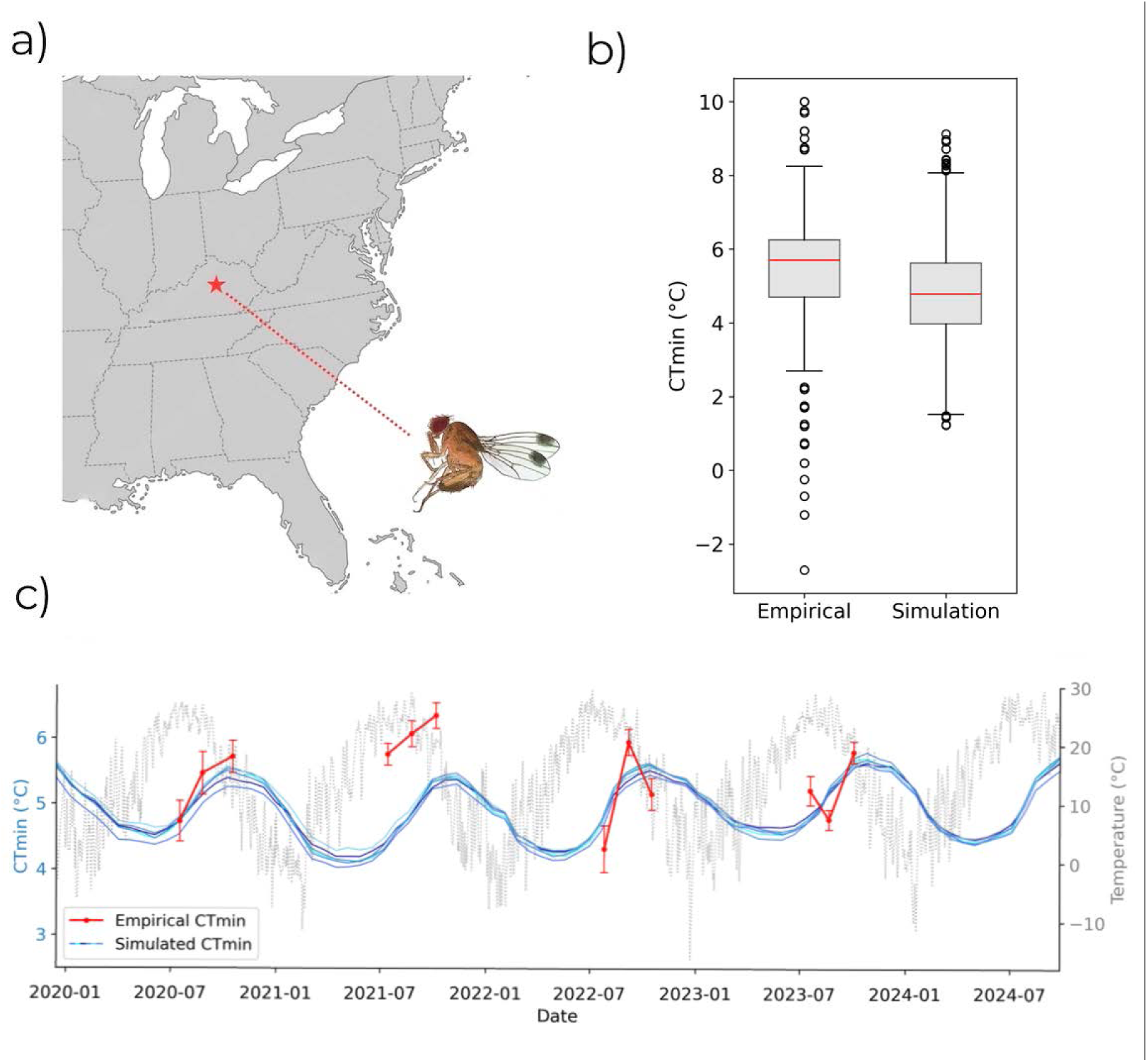
A) Map of the eastern United States showing the *Drosophila suzukii* empirical collection site in Berea, Kentucky (red star). B) Box plots of individual *CTmin* variability comparing empirical measurements (n = 1741) to simulated individuals (n = 1741) from the generations experiencing the equivalent body temperature. C) Seasonally sampled empirical *CTmin* closely aligns with simulated *CTmin*, with both tracking the underlying body temperature distribution. The gray dotted line shows the empirical daily temperature time series (right y-axis) experienced by both empirical and simulated populations. Red points indicate empirical mean *CTmin* measurements from seasonally collected populations, with error bars representing ± 2 SE, and connecting red lines denoting a single season. Colored lines show 5 simulation replicates parameterized with *B*_extreme_ of 30*^◦^*C, *CTmin*_extreme_ of 1.5*^◦^*C, a generation length of 21 days, a fixed population size of 5000 individuals, and a quantitative trait nucleotide mutation rate of 1 × 10*^−^*^6^ per base pair with each mutation equally likely to affect either *B* or *CTmin*. Other parameters were given following the Methods.

To test if our model could capture this empirical pattern, we input empirical temperature data spanning from 1981-01-01 through 2025-01-01 for the collection site coordinates (37.5915°N, 84.3077°W; Figure 4A) as proxy for body temperature, obtained from NASA’s Prediction of Worldwide Energy Resources (POWER) dataset. We fit the simulated *CTmin* to the empirical *CTmin* by manual adjustment of population size, the quantitative trait nucleotide mutation rate, the variance in mutational effects, the generation length in days, and physiological constraints for *B* and *CTmin* (see Methods).

In three of the four sampling years, replicated simulations were able to accurately model the mean *CTmin* observed in the empirical data (Figure 4C). In addition, our model captured the multi-generational lags observed in all years (Figure 4C). We also tested if our model captured the individual variability in *CTmin* observed in the empirical data by sampling the same number of individuals at each sampling event. Across all sampling events (*n*_flies_ = 1741), the distribution of *CTmin* values from the simulations was similar to those empirically observed (empirical mean 5.45 and standard deviation 1.48; simulation mean 5.69 and standard deviation 1.16), with the exception that the empirical dataset had a few individuals with more extreme *CTmin* (Figure 4B).

#### Other factors affecting adaptive tracking

In this section, we consider two other biologically relevant factors that influence how well a population can track the seasonal body temperature change: temperature-dependent generation time and finite population sizes.

So far, we assumed that there are a constant number of days per generation throughout each simulation regardless of the body temperature at each timestep. However, most ectotherms typically have shortened generation times at higher temperatures [47]. Therefore, we modified the sinusoidal body temperature case study in Figure 3 to model generation length linearly decreasing with body temperature. We compared temperature-dependent generation length to control simulations with fixed generation length that had the same number of generations per year. Variable-generation length simulations had higher annual average *CTmin* with longer lags in winter (Figure S12) compared to controls. This is because there were more generations in summer than winter, and the population adapted faster to increased temperature over the same number of days. Yet, we see clear oscillation in *CTmin* in both cases, meaning the population adaptively tracked selection pressure from both cold and hot temperatures even with a smaller number of generations in winter.

On the other hand, genetic drift can hinder adaptive tracking. We repeated the sinusoidal example from Figure 3 with the population size increased or decreased by an order of magnitude (i.e. *N* = 500 or 50, 000; Figure S13). Simulations with smaller populations has smaller oscillations in *CTmin* than the simulations with bigger populations, and sometimes went extinct (2 simulations out of 30 replicates went extinct for lower *N_e_*). Both medium-sized and larger populations tracked seasonal changes without extinction events, and the temporal lag between body temperature and TPC parameters decreased in larger populations due to less genetic drift (Figure S14).

#### Effect of temperature extremes and autocorrelation on thermal limits

As extreme weather events become more frequent, their influence on the evolution of thermal extremes is an important and open question in predicting population persistence. Although many studies have modeled extreme events by simulating an increased probability of extreme temperatures [15, 48], this approach ignores seasonal autocorrelation in body temperature. The influence of autocorrelation vs. amplitude on the evolution of thermal extremes remains debated [10]. Using real (autocorrelated) daily temperature data with extreme low and high temperatures from Vermont (latitude=-72.47167, longitude=44.36272), we found that the thermal performance curve that evolved in this autocorrelated timeseries had a wider breadth than what is theoretically expected from a bimodally distributed body temperature with the same mean and variance (Figure S15). It was also wider than the scrambled Vermont temperature data, which had the same probability of extreme events but lacked autocorrelation (Figures S15 and S16). The finding that the breadth of TPCs in autocorrelated timeseries evolved to be broader than equivalent series with no autocorrelation is consistent with theoretical models of plasticity evolution. A recent review of evolutionary theory highlighted that more plasticity evolves in environments with higher predictability of fluctuations (i.e. autocorrelation) than in environments with large amplitude of temperature fluctuations [10]. In addition, simulations with autocorrelated temperature evolved lower average critical thermal minimums that were closer physiological limit (*CTmin*_extreme_ = 1.5*^◦^*C), hinting that predictability led to better adaptation to extreme temperatures.

## Discussion

We presented a novel framework for the study of TPC evolution based on an individual-based model grounded in population and quantitative genetics, and is an advancement over previous models because it incorporates physiological limits, allows real time-series as input, and can simulate the complexities of linkage, drift, and genetic architecture. Our novel framework reinforces classical insights into TPC evolution, such as the evolution of thermal specialists in less variable environments and of thermal generalists in more variable environments, but also shows that variation in temperature cannot always predict the breadth of TPCs and explains why species may have thermal performance curves that do not match their temperature exposure. We showed that our framework can be used to model adaptive tracking and accurately capture seasonal patterns of *CTmin* in a short-lived insect. The model will have many applications in biological research, that range from giving insight into observed empirical patterns, to creating hypotheses that could be tested experimentally, to predicting species responses to climate change.

### Considerations regarding Jensen’s inequality

Jensen’s inequality states that thermal performance at average temperature is not equal to the average performance in fluctuating environments, with the latter being more accurate [18, 42, 48]. Our model accounts for the effect of Jensen’s inequality by defining lifetime fitness as the average of each timestep’s fitness (conditional on their lifetime history, i.e., excluding timesteps after lethal temperature is reached). As a result, we see a clear difference in the evolved TPCs between populations with the same mean temperature but different individual variation in body temperature in both simulation and analytical theory.

A strength of our simulation approach is that it can take any time series of body temperature, with any timestep, as input. Applications of the simulation model, however, may still be susceptible to Jensen’s inequality based on the resolution (i.e., timestep) of the time-series used. Diurnal fluctuations can expose individuals to non-Guassian, fitness-reducing extremes that can influence TPC evolution [28]. Using mean daily temperature as input into the model essentially assumes that daily fitness is equal to the average temperature on that day and is not impacted by hourly or shorter diurnal fluctuations (i.e., not impacted by Jensen’s inequality). On the other hand, brief exposure to extreme temperatures may have minimal effects on fitness. In this study, we found that using mean daily temperature data as a proxy for body temperature was sufficient to predict seasonal adaptive tracking in a short-lived insect. Future studies, however, should carefully consider the timestep used as input to avoid the pitfalls of Jensen’s inequality.

### Insights into TPC evolution

Our results are consistent with previous theories on the effects of thermal variation on TPC evolution, but also offers new insights. Our simulations emphasize the importance of physiological constraints, genetic drift, seasonal fluctuations, and autocorrelation in TPC evolution. We provided quantitative evidence for the theory that predictable (e.g., autocorrelated) fluctuations lead to more plasticity [10], measured here as greater breadth in the TPC. Specifically, we showed that TPCs that evolved in seasonally fluctuating environments (i.e. with autocorrelation) tended to be broader than TPCs that evolved in response to a series of temperatures sampled from the same distribution independently each day (i.e. no autocorrelation). Additionally, our simulations are able to model individual-level variability, which is an advance over previous studies that assume a single TPC for an entire species [48], and we showed that our simulations could predict the variability in a TPC parameter in a wild insect. Our model captures scenarios where thermal optimum evolves to exceed mean body temperature (consistent with previous theory, ref [2]), but provides new insight into scenarios that would cause thermal optimum to evolve to be less than mean body temperature as temperatures exceed the physiological limits. Testing predictions from our model experimentally is a fruitful area for future research.

### Insights into Adaptive Tracking

Early research predicted that alleles with positive effects on thermal breadth would fix in fluctuating environments [23]. This prediction has been challenged with empirical evidence of allele frequencies fluctuating with the seasons [7, 33]. Initially, models that sought to explain seasonally fluctuating allele frequencies assumed seasonally switching dominance (or “segregation lift”) since it helps maintain polymorphisms in fluctuating environments [9, 49]. More recently, others have shown that stable polymorphisms in fluctuating environments can be maintained without reversal of dominance [9, 13, 14]. This body of theory, however, is not useful for understanding the dynamics of seasonally fluctuating quantitative traits for two reasons: they are focused on the maintenance of individual polymorphisms with direct fitness effects, and they made simplifying assumptions of binary switches between “winter” and “summer” [9, 13, 14, 49].

Our model, which uses body temperature timeseries as input, and in which polygenic loci affect fitness indirectly by changing TPC parameters, successfully reproduced adaptive tracking of TPCs without special parameterization like reversal of dominance. While we have not studied the seasonal fluctuations of QTN alleles in our model, the polygenic architecture would likely make antagonistic pleiotropy (alleles being strongly favored in one season and disfavored in the other) weaker for two reasons. Firstly, polygenic architectures are more transient [50]. Secondly, in local adaptation to heterogeneous environments traits can evolve allele frequency clines along a selective cline without similar clinal patterns in the underlying alleles [32], and it is possible the same dynamic could play out through time. Whether traits can adaptively track fluctuating environments without parallel tracking in allele frequencies is an interesting avenue for future research.

Interestingly, we also found that TPC parameters evolve in response to temperature change with a multi-generation lag. The empirical study that we used to compare to our model, which sampled a fully wild population of *D. suzukii*, found long 60-100 day lags between temperature and CTmin [34] (4-8 generations). Most of the evidence for adaptive tracking comes from *Drosophila*, which have short generation lengths relative to seasonal temperature fluctuations. Previous empirical studies in *Drosophila* have reported a lag of about 14 days (1̃ generation) beteween temperature and an evolutionary response. However, many of these studies were conducted either in large outdoor cages, which may alter certain evolutionary drivers (i.e., potentially limited gene flow/migration, population size changes, mating availability, etc.) [34, 41], or, they have been defined relative to an arbitrary temperature threshold rather than the mean temperature [33], or from patterns restricted to specific loci [36]. The model of TPC evolution presented in this study accurately captured these multi-generational lags in *D. suzukii* in the wild, and suggests that the delay between the action of selection and its response may be much longer in the wild than previously assumed. These multi-generational lags can be explained by the accumulation of allelic effects of highly polygenic traits [29]. In addition, our model predicts that lags between temperature and *CTmin* will be longer coming out of winter than coming out of summer, which is due to the asymmetric shape of the TPC with steeper fitness declines with warming and the higher variance in fitness in summer. This prediction could tested with year-round sampling.

### Limitations and Future Directions

With a flexible simulation framework in hand, how can this model be applied to a specific species? One challenge in applying our modeling framework to a specific species is the requirement of a good estimate for body temperature, which often is quite different from ambient temperature because it can be influenced by microclimate and behavior [46]. Existing biophysical models that estimate microclimate experienced by organisms based on physiological and environmental factors could be a useful way to get proper input temperature data for our simulation [35]. In addition, empirical estimates of physiological extremes are also required as input to the model. For example, in our application to *D. suzukii*, we used the 0.01 quantile of empirically observed *CTmin* as the physiological extreme in the *w_CT min_* function. Finally, empirical data on the genetic architecture of the TPC parameters in a specific species would be useful for parameterizing the mutation rate underlying the QTNs in the model. Even if some of these biological details are missing, it is still possible to generate hypotheses to test, as discussed above.

The persistence of species under such thermal stress depends on their ability to respond through mechanisms such as phenotypic plasticity, behavioral thermal regulation, and evolutionary adaptation [3]. Although integrating the non-genetic mechanisms will be an important step to improve our simulation, our simulations are nonetheless an improvement over current models that estimate extinction risk and range shifts due to global climate change based on a fixed TPC per species [5]. A next natural step would be to address adaptation to other environmental stressors co-varying with temperature, such as insecticides [38], and to implement more realistic demographic pattern including boom-and-bust patterns [27, 34, 36] and migrations across different climate zones (note ref [9] found boom-and-bust to contribute to adaptive tracking in a single-locus model). Individual-based quantitative genetic simulations, like that presented here, will increase the accuracy of predictions of extinction risk, loss of biodiversity, and change in species distributions due to climate change.

## Materials and Methods

### Data availability

All scripts and workflow to reproduce our simulation results are available on https://github.com/jiseonmin/TPC_evolution_SLiM and will be archived with Zenodo upon acceptance.

### Analytical Model: constructing the fitness landscape

#### Thermal Performance Curve (TPC) Parameterization

While there is variation among species due to differences in behavior, physiology, and habitat, typically thermal performance is thought to rise exponentially from a low body temperature due to the thermodynamics of enzymatic activities, peak at an optimal body temperature (*Topt*), and fall rapidly as body temperature gets close to the critical thermal maximum (*CTmax*).

The breadth (*B*) is the difference between *CTmax* and *CTmin*. A typical TPC is described mathematically as a left-skewed unimodal distribution that is zero higher than *CTmax* and exponentially decays toward 0 as body temperature approaches the critical thermal minimum (*CTmin*). Following this conventional knowledge, we chose a half-Gaussian-half-parabolic TPC model from Deutsch et al. [19] as a baseline for our fitness model. Additionally, we consider *CTmin* and *B* as quantitative traits while constraining *Topt* = *CTmin* + 2*B/*3 and therefore keeping the number of free parameters to two for simplicity. With this, the enzymatic fitness component *w*_enzymatic_ is

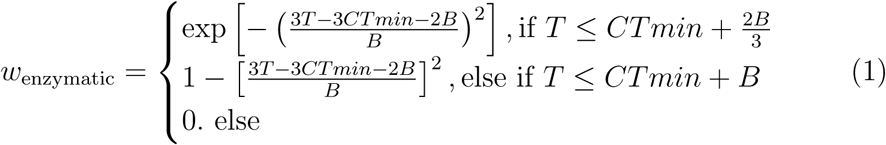

#### Fitness costs for expressing physiological extremes

A novel aspect of our model is the incorporation of a fitness costs for expressing extreme values of *CTmax* (i.e. extreme heat tolerance), *CTmin* (i.e. extreme cold tolerance), and *B* (extremely broad temperature range). In other words, fitness declines as these parameters become more extreme. We define the thermal performance curve *w_T PC_* as a product of four fitness components:

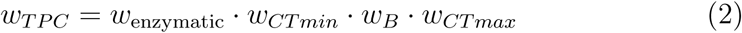

The first component *w*_enzymatic_ is defined in Eq. (1). Each of the other three fitness components is modeled as a logistic function, with Δ controlling how gradual fitness component changes between 0 to 1 around the physiological extreme (see Figure 1 and Figure S1 for visualization). We define *B*_extreme_, *CTmin*_extreme_, and *CTmax*_extreme_ that incur a relative fitness cost of 0.5 (50% reduction in fitness).

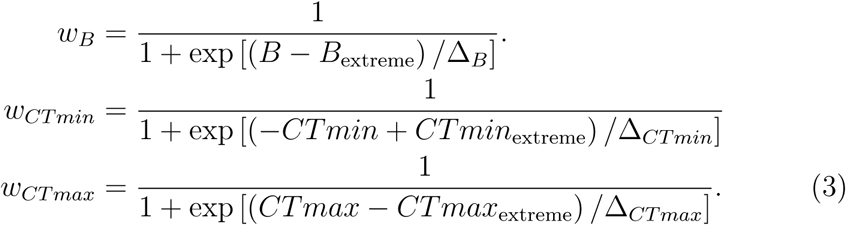

#### Fitness landscape from fitness components in fluctuating temperature environments

In a constant temperature condition, a fitness landscape is simply the same lifetime-fitness function (Eq. (2)), but plotted in a two-dimensional surface with respect to various *CTmin* and *B* values while keeping *T* a constant (Figure S2).

To incorporate fluctuations in body temperature within an individual’s lifetime, we use the expected fitness landscape, *E*[*w*_lifetime_], which is obtained by averaging the fitness landscape *w*_lifetime_ over a probability distribution of temperature *T*, *p*(*T*).

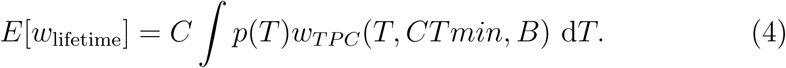

In SI Method A we derive and describe how the coefficient *C* in Eq. (4) accounts for lethal temperatures (e.g., the organism has 0 fitness after temperature exceeds *CTmax* as shown in Figure 1). In the SI we also show how to a gradient of Eq. (4) predicts the evolutionary trajectory for a given TPC (Figure 1 Method 1).

### Simulation model

We use SLiM 5.0 to simulate a forward-in-time Wright-Fisher model of a diploid population with a constant population size, which is 5000 by default unless otherwise noted. The default values for simulations were *B*_extreme_ = 40, Δ*_B_* = 2*, CTmin*_extreme_ = 0, Δ*_CT min_* = 2*, CTmax*_extreme_ = 40, Δ*_CT max_* = 0.2 (see Eq. (3)). In the simulations, fitness of each individual was determined by genotype, phenotypic noise and body temperature.

#### Genetic Architecture & TPC determination

Across all simulations, each individual carried a chromosome containing a 60 kb long QTN region flanked by two neutral arms, each 20 kb long. The chromosome was divided into 12 linkage groups, each containing 5 kb of QTN region, and the first and last with the neutral region attached to the left and the right. We used a uniform recombination map where the rate of recombination was 1e-8 per base pair per generation.

In our simulation model, *CTmin* and *B* were modeled as quantitative traits whose values are determined by the mutations in a quantitative trait nucleotide (QTN) region and phenotypic noise. We assumed that there were two types of QTN mutations, one with additive effects on *CTmin* and the other on *B*, with a mutation rate of 1e-7 per base pair per generation by default and an chance of a new mutation affecting one of the traits. In the default parameter set, the effect size was drawn from a normal distribution with zero mean and variance 0.05. An individual without any QTN mutations had a TPC parameterized by *B*_0_ and *CTmin*_0_. By default, simulations had the initial value of *B*_0_ = 31 and *CTmin*_0_ = 5 (but see Figure S4 for alternative initial conditions). As individuals accumulate QTN mutations, *B* and *CTmin* are calculated as the sum of the effect size of their respective QTNs:

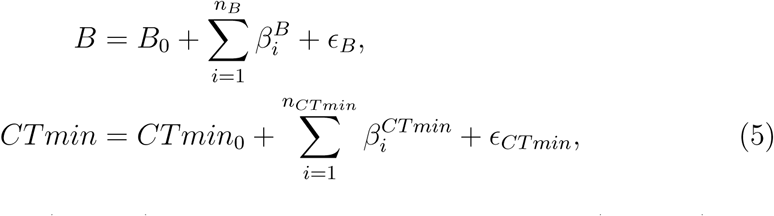

where we have *n_B_* (*n_CT min_*) number of QTN mutations for *B* (*CTmin*), and 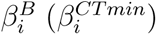 is the effect size of the *i*-th mutation. In addition to the genetic factor, the phenotypes *B* and *CTmin* are influenced by environmental noise, *ɛ_B_* and *ɛ_CT min_* that are sampled from a normal distribution with zero mean and default standard deviation 0.5.

#### Body temperature as a function of time

In the simulations we specify the body temperature that an individual experiences at each timestep as input to the fitness function. By default individual body temperature equaled the population-mean temperature at a timestep plus a random Gaussian noise with a standard deviation *σ_T_*. In the first part of our analysis (Figure 2), we assumed a constant temperature (*µ_T_*). In the second part of our analysis (Figure 3, Figure 4), we modeled mean body temperature of the *i*-th timestep using an external time-series data, which can be synthetic like a sine wave or empirically measured (NASA POWER data). For each individual body temperature on each day we added an independently sampled Gaussian noise with standard deviation *σ_T_* = 1*^◦^*C (Figure 3 and Figure 4).

The default values in the simulations assumed each generation lasted for 10 timesteps (i.e. number of timesteps per generation *n_r_* = 10 and *i* = 1, 2, · · · 10), if not specified otherwise. The fitness of the individual is calculated from the TPC function for a body temperature or is 0 for all the timesteps after a lethal thermal event happened, averaged over all timesteps in their lifetime (see Eq. (S7)).

#### Other simulation details

Every simulation was repeated 30 times with a unique random seed to ensure reproducibility. Each simulation included a burn-in period (5k generation) where all individuals in the population had equal fitness regardless of their genotype or temperature. This burn-in period was added to create standing genetic and phenotypic diversity before selection was implemented. The runtime was initially set to 20k generations for all simulations and extended as needed until equilibrium was reached.

For the simulations using empirical temperature data from Kentucky (Figure 4), we looped through the 1981-2025 temperature time series 20 times in order to model more generations than data was available for. For simulations used to compare with the empirical data, we used the following parameters: population size *N_e_* = 5, 000; mutation rate per base pair = 1×10*^−^*^6^ [1]; generation length = 21 days; small additional temperature variation between individuals (stdev = 1*^◦^*C); *B*_0_ = 30*^◦^*C; *CTmin*_0_ = 0*^◦^*C; *B*_extreme_ = 30*^◦^*C; *CTmin*_extreme_ = 1.5*^◦^*C [17, 26, 43]; *CTmax*_extreme_ = 40*^◦^*C [37, 39, 43]. We sampled simulated individuals on dates that aligned with the empirical sampling (for adaptive tracking *n* = 5000 and for individual variability *n* equaled the empirical sample size).

For auto-correlated simulations in Figure S15, we used NASA POWER daily temperature in Vermont (Longitude = −72.47167, Latitude = 44.36272) from 2015-1-1 to 2020-12-31, looping through the data 200 times. For the non-autocorrelated case, body temperature was randomly sampled from the timeseries everyday. The same parameter values as the Kentucky case study were used in both autocorrelated and non-autocorrelated simulations.

### *D. suzukii* data collection

We sampled populations of *D. suzukii* three times per year in the summer (about six weeks apart), for four consecutive years at two farms in central Kentucky by collecting infested berries and allowing flies to emerge. We quantified *CTmin* in both the parental (P) generation and in the F4 generation of isofemale lines derived from these collections. The fruit containing the P generation was kept at room temperature (20*^◦^*C), and emerging adults and subsequent lines were maintained at 25*^◦^*C until measurement of *CTmin*. *CTmin* measurements in the F4 generation were used to compare to the simulations in this study, and were conducted on flies from up to 20 isofemale lines from each collection. See full details in ref [34].

## Acknowledgments

This work was supported by National Science Foundation Grant 2412803 to KEL, JCBN, and NT. ZC was additionally supported by a PEAK Summit Award from Northeastern University. We thank the Lotterhos lab, Mark Bitter, and Lauren Buckley for their feedback on the manuscript, and Northeastern Research Computing for computing resources and support.

