## Supplementary material for "A general model for the evolution of thermal performance curves with application to real time-series data": SI

### 1 SI Methods

#### 1.1 Effect of a lethal thermal event

In our fitness model, we assume that the fitness of an individual comes from the arithmetic average of reproductive output per timestep. Note that prior literature discovered how Jensen's inequality (i.e. mean of fitness over temperature distribution is higher/lower than fitness at mean temperature if TPC is concave down/up around the temperature distribution) implies that accounting for variance of temperature (not just average) is necessary for accurately predicting thermal performance [3]. Once the individual is exposed to lethal body temperature, it stops reproducing, and even if the body temperature in the following timesteps returns to a sub-lethal level, there is no further reproductive output contributing to the fitness. We will call this the "no-recovery" model, to distinguish it from the "recovery model" where daily reproductive output solely depends on the body temperature of that timestep.

###### 1.1.1 Recovery model

In this simpler "recovery model", expected lifetime fitness is simply

$$E[w_{TPC}]_{\text{recovery}} = \int p(T) w_{TPC}(T) dT, \tag{S1}$$

(i.e. TPC averaged over the probability density function of body temperature). This is equivalent to the fitness measure in previous studies on effects of thermal fluctuations and Jensen's inequality (e.g. [4]), but it is not able to account for time-dependent nature of thermal damage. In other words, how early or late temperature goes outside the viable range does not affect the total reproductive output, implying that organisms can completely and immediately *recover* from thermal damage.

###### 1.1.2 No-recovery model

The derivation of the no-recovery model requires a definition of a viable range. The viable range can be either defined as the range between $CTmin$ and $CTmax$ ($CTmin < T < CTmax$) (symmetric effect) or it can also just have the upper limit ($T < CTmax$) if exceeding $CTmax$ causes mortality but temperatures below $CTmin$ do not (asymmetric effect).

Let's say the temperature $T$ gets outside the viable range for the first time at the $c$-th timestep. Then the fitness at $i$-th timestep is zero if $i \geq c$. And before the $c$-th timestep, the expected fitness should reflect the average over temperature within the viable range only. So expected fitness at the $i$-th timestep, conditional on that the organism gets outside the viable thermal range for the first time on the $c$-th timestep is

$$E[w_{\text{timestep}}(i)|c] = I(i-c)\frac{1}{r}\int p(T) w_{TPC}(T) dT \tag{S2}$$

where $r$ is area under the normal distribution $p(T)$ truncated to only include the viable temperature range:

$$r = \int_{-\infty}^{CTmax} p(T)dT,$$

if only overheating has irrecoverable damage (asymmetric), or

$$r = \int_{CTmin}^{CTmax} p(T)dT,$$

for symmetric effect from overheating and freezing.

To find the expected lifetime fitness for a given $c$, we should average the expected fitness (Eq. (S2)) over all $n_r$ timesteps (assuming there are $n_r$ timesteps per generation).

$$E\left[w_{\text{lifetime}}|c\right] = \frac{1}{n_r}\sum_{i=1}^{n_r} E[w_{\text{timestep}}(i)|c] = \frac{c-1}{rn_r}\int p(T)w_{TPC}(T)dT. \quad \text{(S3)}$$

To find the expected lifetime fitness $E\left[w_{\text{lifetime}}\right]$, we need to consider all possible values of $c$. Let's call the probability that $c = j$ (i.e. first thermal lethal event happens on the $j$-th timestep), $P(c = j)$. Note that a lethal event may not happen at all. We will call that possible scenario without thermal lethal event "$c = n_r + 1$" and its probability $P(c = n_r + 1)$, since it will be convenient for summation notation later. The conditional mean fitness $E[w_{\text{lifetime}}|c = n_r + 1]$ is equal to the mean fitness in the recovery model from before (Eq. (S1)):

$$E[w_{\text{lifetime}}|c = n_r + 1] = \int p(T)w_{TPC}(T)dT$$

This is also consistent with Eq. (S3) and shows that Eq. (S3) generalizes to $c = 1, \cdots, n_r + 1$.

So combining all this, we find a simpler equation of the expected lifetime fitness:

$$\begin{aligned} E\left[w_{\text{lifetime}}\right] &= \sum_{j=1}^{n_r+1} P(c = j)E\left[w_{\text{lifetime}}|c = j\right] \\ &= \sum_{j=1}^{n_r+1} P(c = j)\frac{j-1}{rn_r}\int p(T)w_{TPC}(T)dT. \end{aligned} \quad \text{(S4)}$$

Lastly, we need to know what $P(c=j)$ is to calculate $E\left[w_{\text{lifetime}}\right]$. To do so, we will use $r$ again, which is the probability of drawing a temperature within the viable range. If $c=j$ and $j<n_r+1$, the first j-1 temperatures are inside the viable range and the $j$-th temperature is outside the viable range. So

$$P(c=j)=r^{j-1}(1-r).$$

If $c=n_r+1$, all $n_r$ independently drawn temperatures are within the viable range.

$$P(c=n_r+1)=r^{n_r}.$$

Applying these to Eq. (S4),

$$\begin{aligned} E\left[w_{\text{lifetime}}\right] &= \left[\frac{1-r}{rn_r}\sum_{j=1}^{n_r}(j-1)r^{j-1}+r^{n_r-1}\right]\int p(T)w_{TPC}(T)dT. \\ &= \left[\frac{1-n_r\cdot r^{n_r-1}+(-1+n_r)\cdot r^{n_r}}{n_r(1-r)}+r^{n_r-1}\right]\int p(T)w_{TPC}(T)dT. \end{aligned} \tag{S5}$$

We use that $\sum_{j=1}^{n_r}(j-1)r^{j-1}=\frac{r-n_r r^{n_r}+(-1+n_r)r^{n_r+1}}{(1-r)^2}$ to simplify the summation portion of the equation going from the second to last to the last line.

So the coefficient $C$ in Eq. (4) is

$$C=\frac{1-n_r\cdot r^{n_r-1}+(-1+n_r)\cdot r^{n_r}}{n_r(1-r)}+r^{n_r-1} \tag{S6}$$

We see the modified formula of mean lifetime fitness (Eq. (S5)) converge to Eq. (S1), which is the expected lifetime fitness for "recovery" model (i.e. $C=1$, which makes Eq. (S5) equal to Eq. (S1)) in two cases. First, if a measurement is made only once in a generation there is no difference between no-recovery vs. recovery models (we confirm this when we set $n_r=1$ in Eq. (S6), as it becomes one). Second, if the viable temperature range is $(-\infty,\infty)$, $T$ can never be outside the viable range, so again, the no-recovery and recovery models will converge. Mathematically, we can verify this by applying $\lim_{(\text{viable range})\to(-\infty,\infty)} r = 1$ to Eq. (S6), which will make it again be one.

In our Wright-Fisher simulations, each individual's fitness is lifetime fitness, consistent to the definition in the analytical model: the average of TPC function's value the given body temperature or 0 if a lethal thermal event happened on any previous timestep.

$$w_{\text{lifetime}} = \frac{1}{n_r}\sum_{i=1}^{n_r} w_i(t),$$

$$w_i(t) = w_{TPC}(T = T_i(t)) \prod_{j=1}^{i} I_{(viable)}(T_j(t)). \tag{S7}$$

Here $I_{(\text{viable})}(T_j(t))$ is an indicator function which is 1 if the $j$-th body temperature in generation $t$ is within the viable range and is zero otherwise.

#### 1.2 From fitness landscape to evolved TPCs: analytical

We can use the fitness landscape to estimate the trajectory of TPCs evolution. A seminal study by Lande and Arnold [2] illustrated how correlated multivariate traits evolve through quantitative genetics. Specifically, the selection gradient determines the strength and direction of selection, which is approximately the slope of the fitness landscape at the population-mean. In our case, the rate of change of mean $CTmin$ (or $B$) per generation ($d\overline{CTmin}/dt$ or $d\overline{B}/dt$) should be proportional to the selection gradient at the population mean for $CTmin$ (or $B$):

$$\frac{d\overline{CTmin}}{dt} \propto \left.\frac{\partial E[w_{\text{lifetime}}]}{\partial CTmin}\right|_{CTmin=\overline{CTmin},B=\overline{B}}$$

$$\frac{d\overline{B}}{dt} \propto \left.\frac{\partial E[w_{\text{lifetime}}]}{\partial B}\right|_{CTmin=\overline{CTmin},B=\overline{B}}$$

The precise ratio between the adaptation rate and selection gradient depends on mutational supply, genetic drift, and recombination rate.

Let's assume that the mutation rate of QTNs for $B$ ($CTmin$) is $U_B$ ($U_{CTmin}$) and the effect sizes ($s_B$ or $s_{CTmin}$) is sampled from a normal distribution of zero mean and standard deviation of $\sigma_B$ ($\sigma_{CTmin}$). Let's also simplify the problem by assuming that each QTN only affects either $B$ or $CTmin$, but not both (i.e. zero mutational covariance). Then we can write an ordinary differential equation (ODE) for $B$ and $CTmin$ as

$$\frac{d\overline{CTmin}}{dt} = kU_{CTmin}\sigma_{CTmin}s_{CTmin}(\overline{CTmin}, \overline{B}),$$
$$\frac{d\overline{B}}{dt} = kU_B\sigma_B s_B(\overline{CTmin}, \overline{B}).$$

Here $k$ is some constant. It is difficult to know what $k$ is exactly without explicitly simulating the population. However, the trajectory that $CTmin$ and $B$ follows still can be found by solving the ODE above, given the initial trait values. Specifically, the trajectory is described as

$$\overline{CTmin}(\tau) = \overline{CTmin}(0) + \int_0^\tau s_{CTmin}\left(\overline{CTmin}(\tau'), \overline{B}(\tau')\right) d\tau',$$
$$\overline{B}(\tau) = \overline{B}(0) + \frac{U_B}{U_{CTmin}}\frac{\sigma_B}{\sigma_{CTmin}}\int_0^\tau s_B\left(\overline{CTmin}(\tau'), \overline{B}(\tau')\right) d\tau'. \quad \text{(S8)}$$

Here $\tau$ is a rescaled time $\tau = kU_{CTmin}\sigma_{CTmin}t$. Note that this prediction for evolutionary trajectory is only valid when body temperature of each timestep is sampled from the same distribution at every generation (there are no seasonal fluctuations across generations) because the integrands $s_{CTmin}$ and $s_B$ has no direct dependence on $\tau'$ in Eq. (S8). In our simulation model, we define the mutation matrix to be symmetric (i.e. $U_B = U_{CTmin} = 5E-8$ and $\sigma_B^2 = \sigma_{CTmin}^2 = 0.05$), making the ratio between them all 1:

$$\frac{d\overline{CTmin}}{d\tau} = \left.\frac{\partial E[w_{\text{lifetime}}]}{\partial CTmin}\right|_{CTmin=\overline{CTmin}, B=\overline{B}}$$
$$\frac{d\overline{B}}{d\tau} = \left.\frac{\partial E[w_{\text{lifetime}}]}{\partial B}\right|_{CTmin=\overline{CTmin}, B=\overline{B}} \quad \text{(S9)}$$

With a given initial mean $CTmin$ and $B$, $\overline{CTmin}(0)$ and $\overline{B}(0)$, and the ordinary differential equations (S9), we now have an initial value problem whose solution is the expected TPC trajectory, $\left(\overline{CTmin}(\tau), \overline{B}(\tau)\right)$.

We numerically solve this initial value problem for each timestep $\tau$ using `scipy.integrate.solve_ivp` [5] until the trajectory reaches the peak of the fitness landscape (i.e. until both selection gradients become zero to numerical precision limit).

To find the optimal $CTmin$ and $B$, $CTmin_{OPT}$ and $B_{OPT}$ (the values with the maximum predicted fitness) we use the function `scipy.optimize.minimize` [5] to find the peak on the fitness landscape. Specifically, the python function solves for the minimum of the negative of the expected fitness landscape:

$$CTmin_{OPT}, B_{OPT} = \underset{CTmin, B}{\arg\min}[-E[w_{\text{lifetime}}(CTmin, B)]].$$

#### 1.3 Quantifying lag in seasonal adaptive tracking

In Figure 3, annual maximum and minimum TPC parameters and temperature are identified for the last 10 years of the simulations, using `scipy.signal.find_peaks` with minimum distance set to 30 generations (note that each year is 36 generations long, and we want to identify exactly one peak and valley per year). Then the difference between the generation when temperature is maximized and when each TPC parameter is maximized for a given year is defined as "summer lag". Similarly, the difference between the generation when temperature is minimized and when a given TPC parameter is minimized for a given year is "winter lag". Reported lags in main text is the average over 10 years. See `SI_figures_and_tables.py` in `https://github.com/jiseonmin/TPC_evolution_SLiM` for the full script.

#### 1.4 Adaptive tracking for different population sizes

We expected the effect of population size on the rate of adaptation to be more pronounced when the temperature fluctuated across generations, in addition to the within-generation variation. In other words, if the thermal environment fluctuated across generations, larger populations would be able to move toward a new trait optimum in a fewer number of generations (i.e. more effective adaptive tracking) due to weaker genetic drift. To test this prediction, we next simulated TPC evolution using a temperature distribution whose mean ($\mu$) oscillates sinusoidally every 36 generation while standard deviation ($\sigma$) is fixed and smaller than the amplitude of the sinusoidal function: $T_i(t) \sim \text{Normal}(\mu(t), \sigma)$ with $\mu = 20\sin(2\pi t/36) + 20$ and $\sigma = 1$. See Figure S13.

##### 1.5 Bimodal temperature distribution

For the bimodal temperature distribution case (Figure S5), each individual samples temperature from a mixture of two normal distributions with equal heights and widths:

$$\begin{aligned} T_i(t) \sim & 0.5\text{Normal}(\mu_T - \Delta_\mu/2, \sqrt{\sigma_T^2 - \Delta_\mu^2/4}) \\ & + 0.5\text{Normal}(\mu_T + \Delta_\mu/2, \sqrt{\sigma_T^2 - \Delta_\mu^2/4}). \end{aligned}$$

The peaks are separated by $\Delta_\mu$ and the overall distribution has mean $= \mu_T$ and standard deviation $= \sigma_T$

#### 2 SI Results

##### 2.1 Effects of bimodality

We found that increasing bimodality of the temperature distribution did not always make the TPC breadth wider: and in some cases increased biomodality could make the TPC narrower (both in theory and simulation, and both for daily or seasonal fluctuations, Figure S5). This is because, in short, when temperature peaks are separated enough, specializing to one of them can incur higher average performance, compared to generalist whose fitness is suboptimal near either of the peaks. More broadly, this result implies that variance of temperature alone cannot be a predictor for evolutionary trajectory of TPCs: if temperature distribution is bimodal enough and especially if one of the peaks is physiologically costly to adapt to, a better strategy can be to abandon the less favorable thermal regime (e.g., extreme cold in Figure S5).

##### 2.2 Sampling TPCs

In Figure 2, we visualize 40 randomly sampled TPCs (using random seed = 4; see Jupyter notebook in the Github repository for full plotting script) in the last 100 generations in one replicate simulation per parameter set (seed for replicate simulation = 29). The number of TPCs to plot was capped at 40 for visualization purposes, and in Table S3 we report the proportion of TPCs of each category from all individuals sampled from all 30 replicate simulations. In Figure S7, we show the mean and standard deviation of key TPC parameters at the end of simulations. It is notable that no individuals had fitness lowered due to extreme thermal generalism, probably because ranges of temperature fluctuation tested are much lower than the threshold (specifically, $B_{\text{extreme}} = 40$) for which we consider a TPC to be an extreme generalist.

#### 2.3 The evolutionary trajectory of the TPC

In Figure S3, we plot the analytical trajectory compared to the average simulated trajectory of $B$ and $CTmin$ on top of the expected fitness landscape $E[w_{TPC}(CTmin, B)]$ for a given mean and standard deviation of temperatures. In all conditions, the average B and CT min from the replicate simulations (black) follows the theoretical trajectory (red) quite closely, obtained from solving Eq. (S9) numerically. As Figure 2 demonstrates, individual TPCs vary within each population. We also repeat the analysis with a different $B_0$ and $CTmin_0$ (see Figure S4). Note that the three environments (top left, top right, and center left in Figure S4) are missing analytical trajectories (i.e. no red dash line). In the three conditions, numerical solution to Eq. (S9) predicts that the average $B$ and $CTmin$ will stay in their initial value $(B_0, CTmin_0)$ indefinitely because selection gradient there is too shallow. However, in simulations, all 6 populations, including the 3 populations that were predicted to be stuck in the initial conditions, evolved TPCs close to theoretical optimum (black lines in Figure S4). This difference between simulations and analytical predictions highlight the importance of genetic drift and an important reason not to solely rely on analytical theory: our equation (S9) that does not account for genetic drift fails to predict that finite populations are able to explore the fitness landscape without selection and eventually find a path toward the global optimum with sufficient selection gradient.

#### 2.4 Population size and TPC evolutionary trajectory

For a constant temperature environment analogous to tropical (i.e. high mean temperature, low standard deviation) environment as defined in Figure 2, we repeated simulations and with population size of 500, 5000 and 50,000. Initial values of $B$ and $CTmin$ were 23.32 and 16.69. For smallest population size, the simulations were run for 50k generations since the population did not reach equilibrium at 20k generations. The rest of the simulations were all run for 20k generations. Other parameters were the same as Figure 2. As population size increases, genetic and phenotypic correlation between $CTmin$ and $B$ at the end of the simulations become more negative (Figure S8).

#### 2.5 Why is adaptation faster in summer than in winter?

According to Fisher's Fundamental Theorem of natural selection, rate of adaptation is proportional to the additive genetic variance of fitness [1]. Although the QTN mutations in our model does not have a clean additive affects on fitness, as fitness is determined by combined effects of all QTNs on CTmin and B and daily temperature distribution of a given generation, we can look at the total variance of fitness as a proxy for genetic variance of fitness, assuming that the proportion of the genetic variance stays roughly constant over time. In Figure S9, we plot variance of fitness divided by mean fitness from the same simulation in Figure 3. In average, variance is much higher in summer than in winters, which implies faster adaptation in summer and winter, consistent with patterns seen in Figure 3. Then why is fitness higher in summer than in winter? This is due to asymmetric shape of the TPC, where fitness rises gradually from CTmin to Topt and falls much more rapidly from Topt to CTmax. For instance, Figure S10 shows that distribution of fitness at mean temperature 0 is much narrower than at 35 given the same distribution of TPCs (which are the minimum and maximum temperatures in Figure 3). With different minimum and maximum temperatures and different degrees of asymmetry in TPCs may change how different fitness variance is in winter vs. summer, and therefore make the rate of adaptation more or less asymmetric in two seasons.

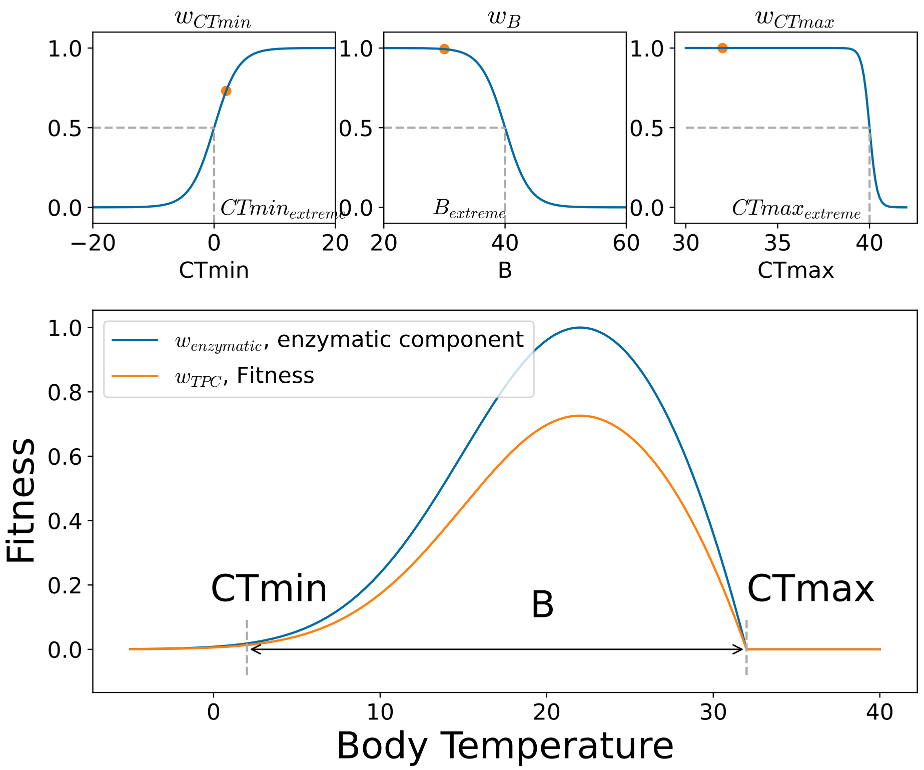


Figure S1: Fitness components as a function of $CTmin$, $B$ and $CTmax$ (top left to right) and $w_{\text{enzymatic}}$ as a function of temperature (blue curve). The orange points mark the values of $CTmin$, $B$ and $CTmax$ from the bottom panel–in this figure, we use $B = 30$ and $CTmin = 2$. The fitness, $w_{TPC}$, is lower than the enzymatic component $w_{\text{enzymatic}}$ by a factor of $w_{CTmin} \cdot w_B \cdot w_{CTmax}$. In this figure, we use $B_{\text{extreme}} = 40, \Delta_B = 2, CTmin_{\text{extreme}} = 0, \Delta_{CTmin} = 2, CTmax_{\text{extreme}} = 40, \Delta_{CTmax} = 0.2$. The gray dash lines marks where the extreme values of trait values, where the fitness component is 0.5 and is declining or increasing at the fastest rate.

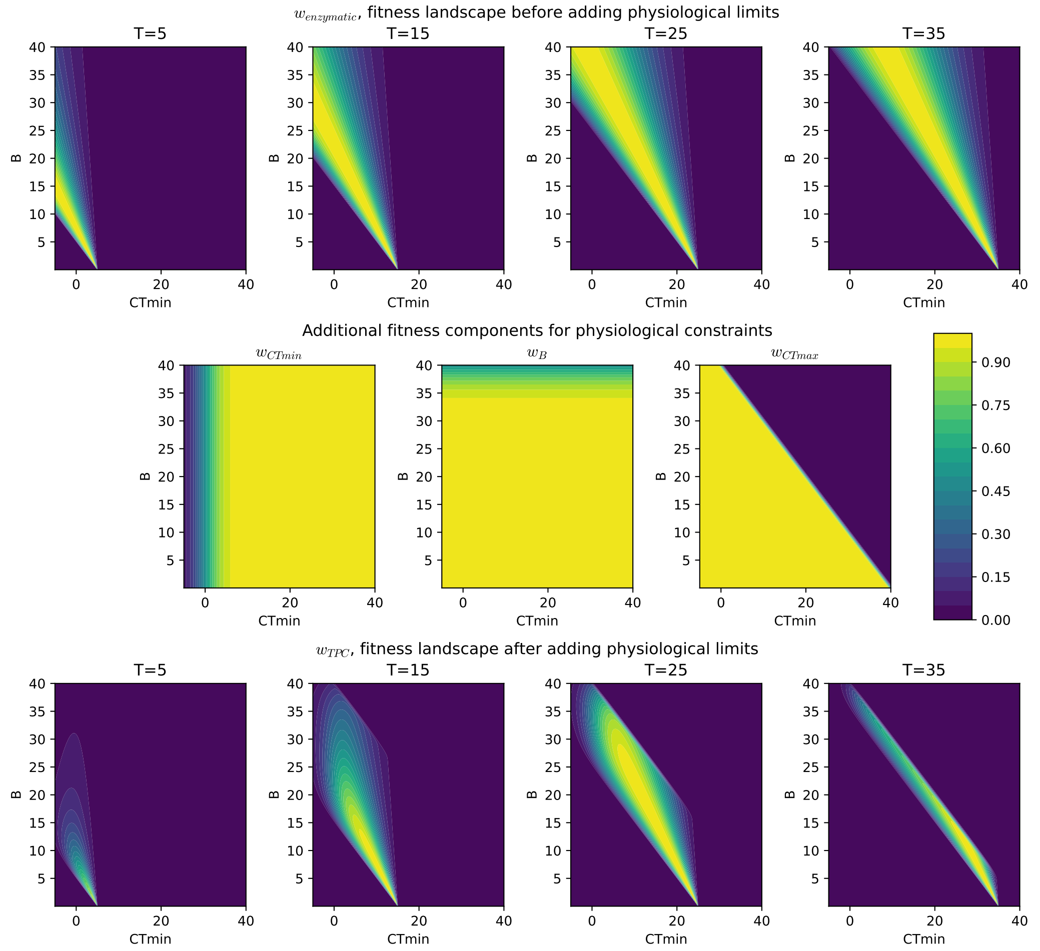


Figure S2: Fitness landscape at a constant temperature before and after adding physiological constraints. Here we set $B_{\text{extreme}} = 40, \Delta_B = 2, CTmin_{\text{extreme}} = 0, \Delta_{CTmin} = 2, CTmax_{\text{extreme}} = 40, \Delta_{CTmax} = 0.2$.

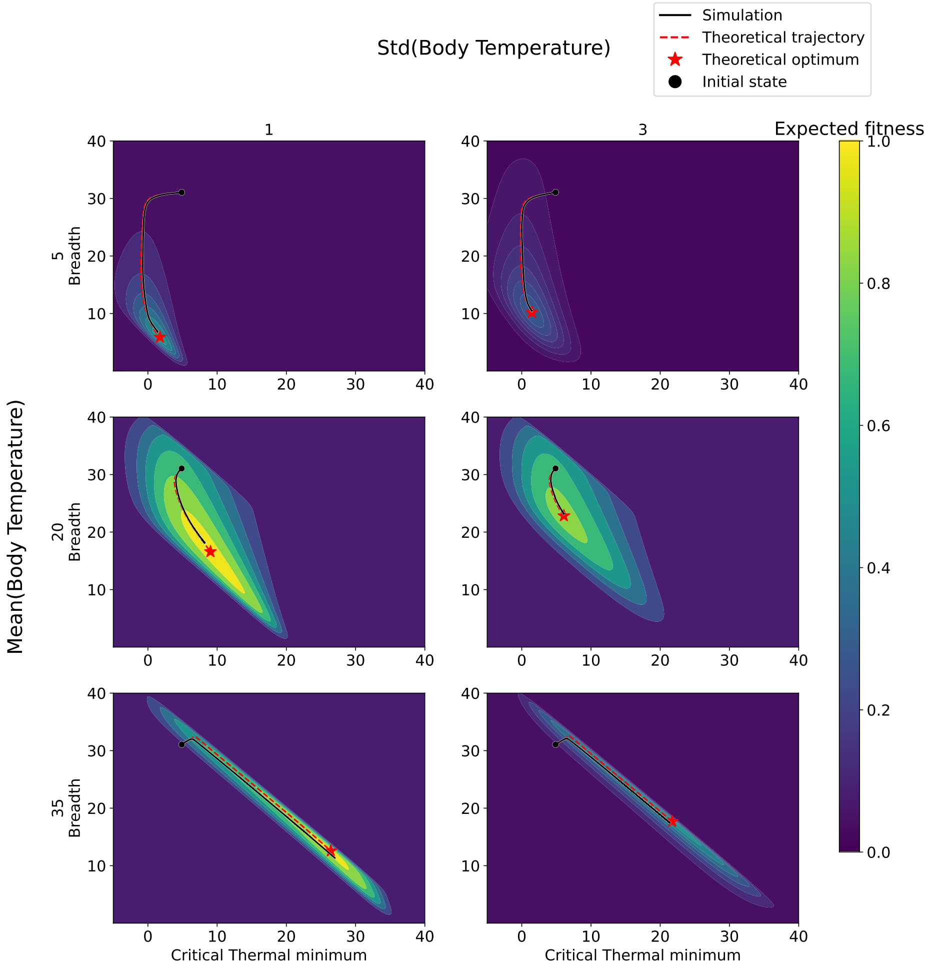


Figure S3: Theoretical TPC trajectory is comparable to the average simulated trajectory. The trajectories are plotted on top of the expected fitness landscape, $E[w_{\text{lifetime}}]$ for a given temperature distribution. The global maximum of expected fitness is found separately (see SI-Methods) and are marked as red stars. In all 6 populations, we set initial traits to $B_0 = 31$ and $CTmin_0 = 5$, marked as black circles. Other parameters are default as described in the main text.

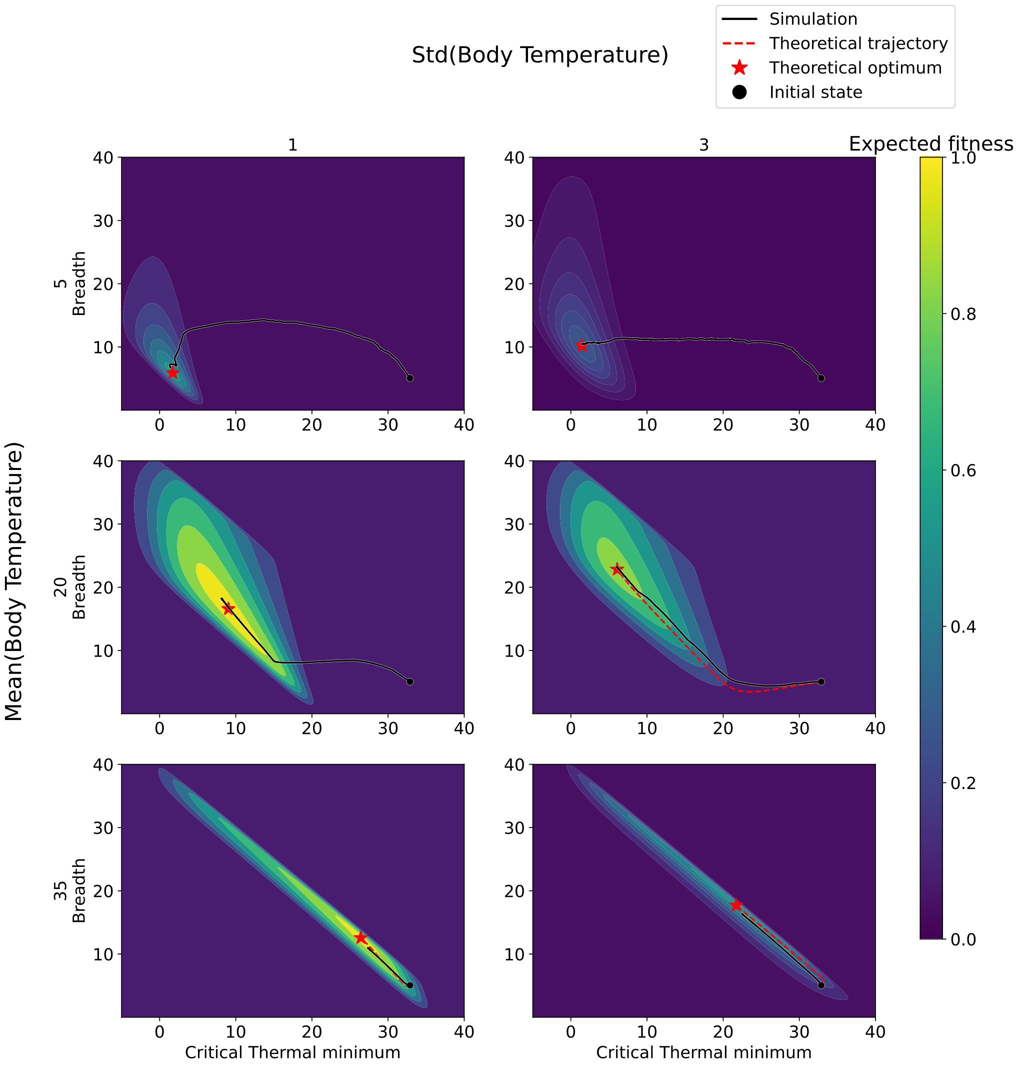


Figure S4: Average TPC trajectories, simulated and theoretical, with different initial conditions compared to Figure S3 – $B_0 = 5$ and $CTmin_0 = 33$. In three conditions where the initial selection gradient is extremely close to zero (top left, top right, center left), the solution to differential equations predicted the TPCs to stay in the initial condition. In simulations under these same conditions, however, where mutations and genetic drift introduces stochasticity to the trait distribution, the population is able to explore the trait space even without a strong selective gradient and is able to find the optimal trait values that maximize the expected fitness $E[w_{\text{lifetime}}]$ (red stars). Other parameters are default as described in the main text.

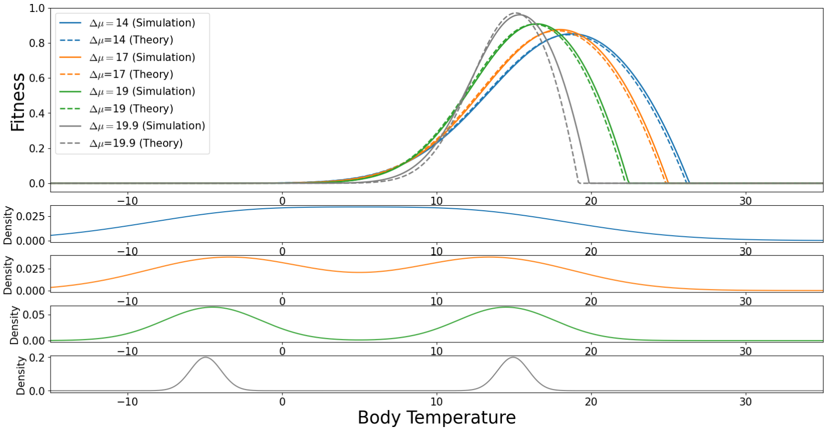


Figure S5: Bimodality can decrease the breadth of TPCs – verified through simulations and theory. Daily temperature is sampled from mixture of two Gaussian distributions with equal height and equal variance ($\sigma^2 - \Delta\mu^2/4$). The means are $\mu - \Delta\mu/2$ and $\mu + \Delta\mu/2$, making total variance $\sigma^2$ and total mean $\mu$, for any amount of separation between the two peaks ($\Delta\mu$). We set $\mu = 5, \sigma = 10$ for all 4 temperature distributions, which is comparable to the Vermont daily temperature data used in Figure S15. Top – Population average TPC from simulations and theoretically optimum TPC. Bottom – Probability density function of daily temperature for given $\Delta\mu$. Parameters are default as described in the main text, with $CTmin_{extreme} = 0$ and $B_{extreme} = 40$.

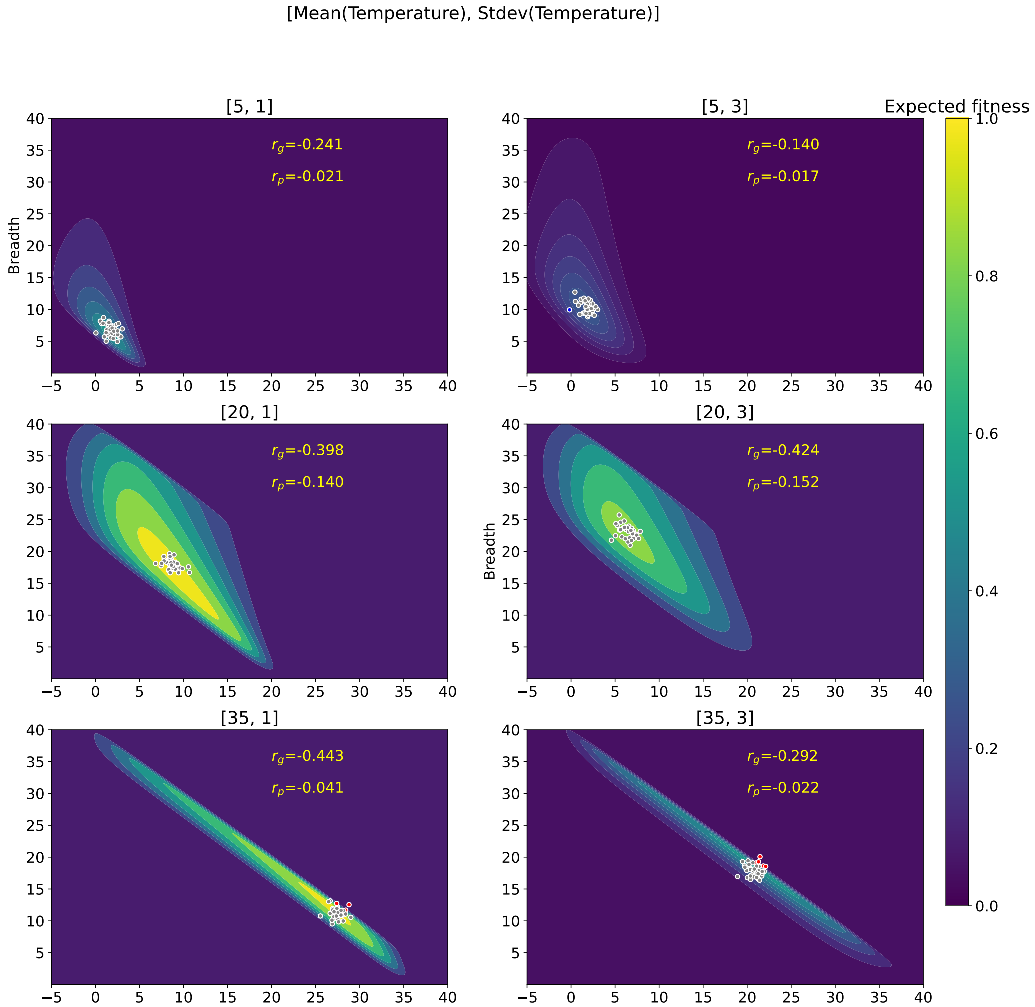


Figure S6: Sample TPCs on expected fitness landscape for all 6 populations shown in Figure 2(a). $r_g$ is genetic correlation between $CTmin$ and $B$ (i.e. correlation coefficient between $\sum \beta_{CTmin}$ and $\sum \beta_B$) and $r_p$ is the phenotypic correlation between $CTmin$ and $B$ (which includes environmental noise). In all 9 populations, genetic correlation ($r_g$) evolves to be negative due to the negative correlation in selective gradient, as shown by the diagonal shape of fitness landscapes. The phenotypic correlation ($r_p$) is less negative than genetic correlation ($r_g$) since environmental noise that was to the trait values in the model.

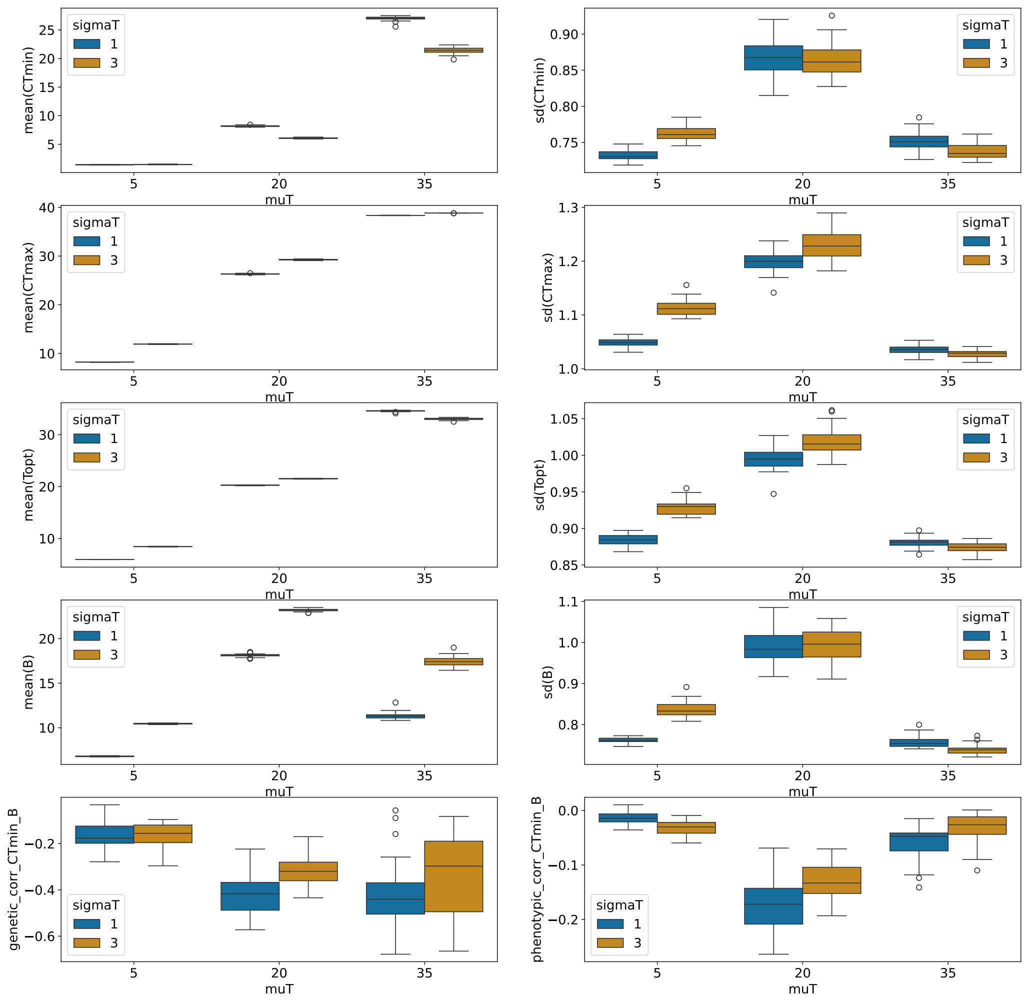


Figure S7: Box plots of final $TPC$ parameters at the end of all 30 replicate simulations used for Figure 2. MuT stands for mean doby temperature, and sigmaT stands for standard deviation of body temperature in constant environments. 8 plots of the first three rows report distribution of mean and standard deviation of TPC parameters of 10,000 randomly sampled individuals (100 per generation for the last 100 generations) in 30 replicate simulations. The bottom right plot reports the distribution of the correlation of $B$ and $CTmin$ from the sampled TPCs. The bottom left plot shows genetic correlation of $CTmin$ and $B$, measured from the effect sizes of segregating QTNs in all individuals at the end of each replicate simulation from its tree sequence output. See Table S1 for more information about the segregating QTNs for CTmin and B.

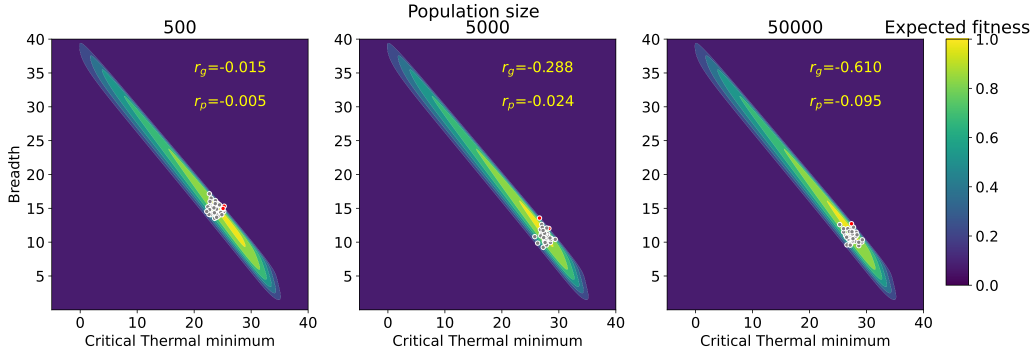


Figure S8: Sample TPCs for "tropical" population (as defined in Figure 2) of various population sizes. $r_g$ is the genetic correlation between $CTmin$ and $B$ (i.e. the correlation coefficient between $\sum \beta_{CTmin}$ and $\sum \beta_B$), and $r_p$ is phenotypic correlation between $B$ and $CTmin$ (which equals the sum of mutational effects plus environmental noise). Drift is weaker in a larger population, so genetic and phenotypic correlations become more negative, as they align better with the negatively correlated selective gradient.

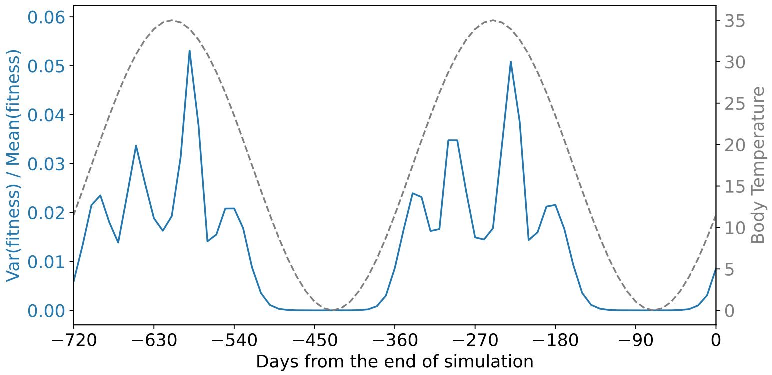


Figure S9: Variance of fitness, normalized by mean fitness, is higher in summer than in winter. Assuming similar additive genetic variance of fitness follows the similar pattern, this is consistent with faster adaptation in summer than in winter in Figure 3 according to Fisher's Fundamental Theorem of Natural Selection [1].

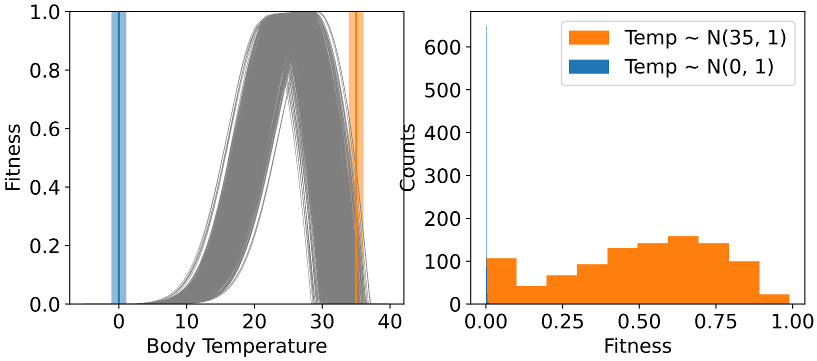


Figure S10: Distribution of daily temperature tends to be narrower in winter than in summer for left-skewed TPCs. Given the same individual TPCs to sample from (grey curves in left), daily fitness of one random individual is measured first for 1000 daily temperatures first sampled from a normal distribution with mean 35 and standard deviation 1 (orange), and second from a normal distribution with mean 0 and standard deviation 1 (blue). The measured daily temperatures is much narrower in winter than in summer (blue vs. orange in the right panel).

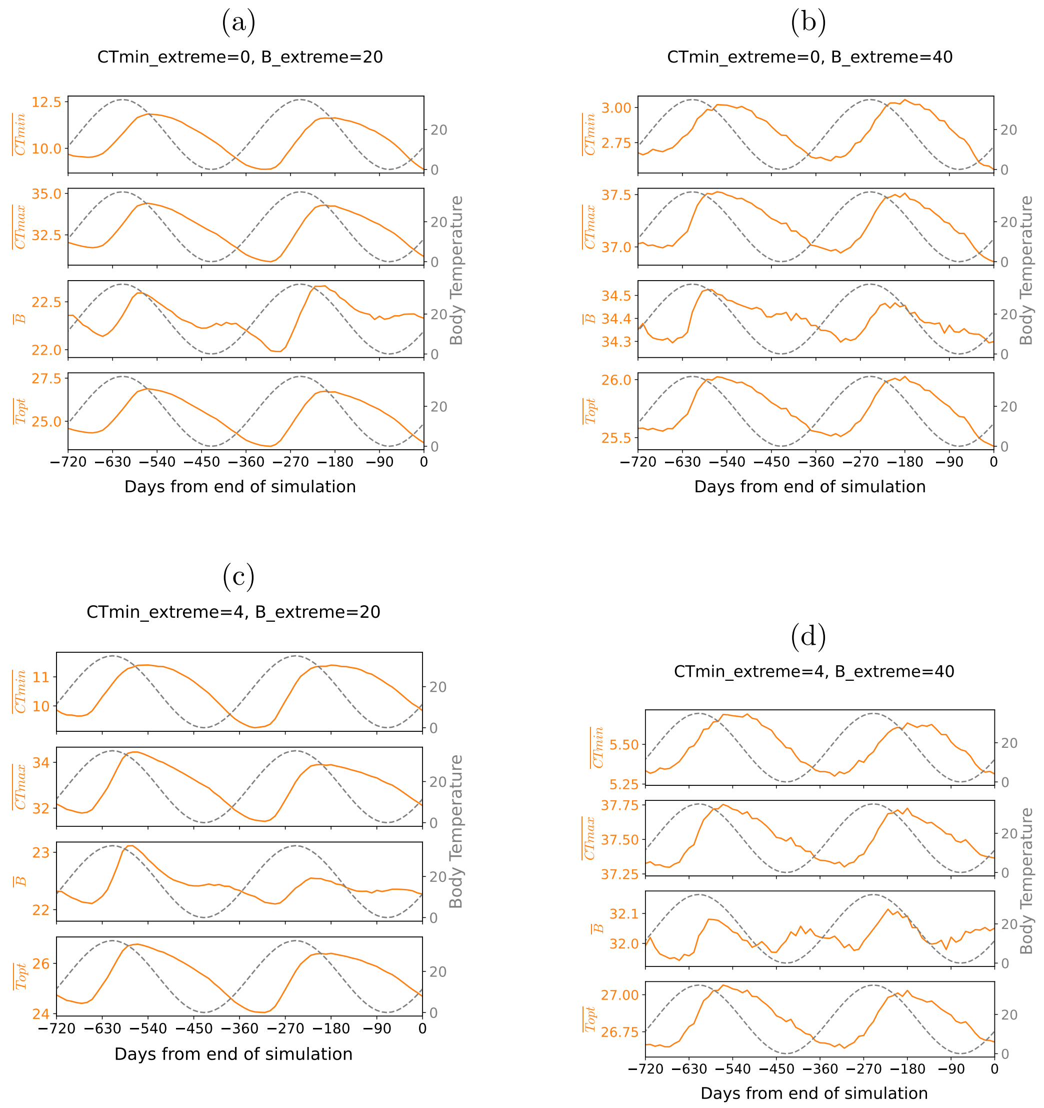


Figure S11: Four key parameters adaptively tracks sinusoidally fluctuating temperature (between 0 and 35 with 360 days period) for all 4 combinations of $B_{\text{extreme}}$ and $CTmin_{\text{extreme}}$. But when $B_{\text{extreme}}$ is smaller ((a), (c) vs. (b), (d)) tracking is stronger (i.e. bigger range of oscillation in all TPC parameters) because generalists are negatively selected.

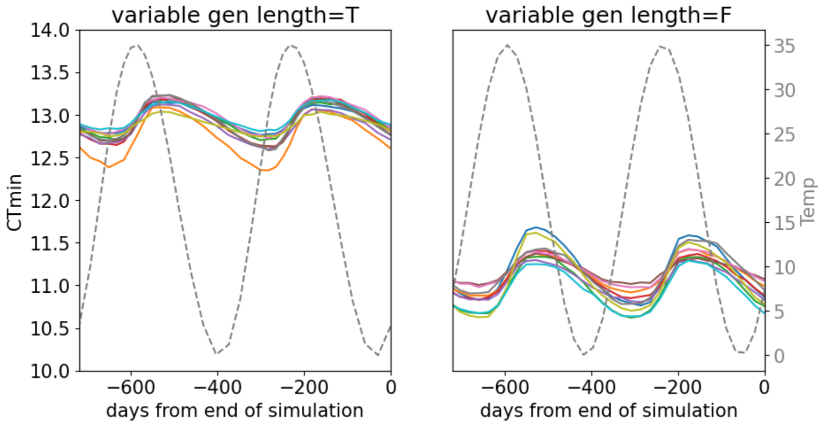


Figure S12: Effects of temperature-dependent generation length. Keeping the annual average the same, generation length is either fixed or made temperature-dependent: specifically, it decreases linearly with temperature between $T = 0$ and 40 from 30 days to 14 days, and stays constant at 30 days for $T < 0$ and 14 days for $T > 40$. As a result of there being more generations in summer vs. winter, population with variable generation length is better adapted to hotter temperature than the one with fixed generation length: higher annual average $CTmin$, faster adaptation to summer, and slower adaptation to winter.

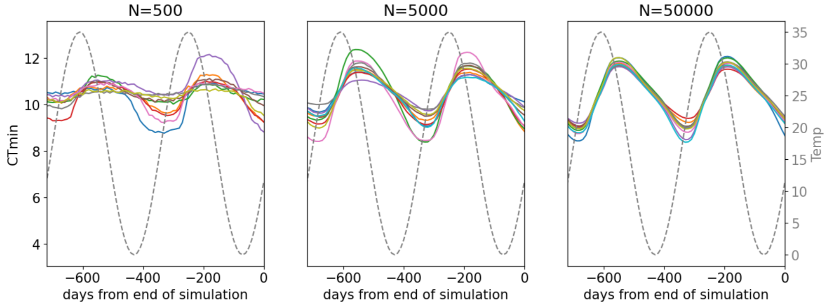


Figure S13: Oscillation of mean $CTmin$ for various population sizes. Each line represents result from random seed = 0 to 9. Simulation with $N = 500$ and seed = 6 went extinct shortly after the burn-in period, so it is not plotted here.

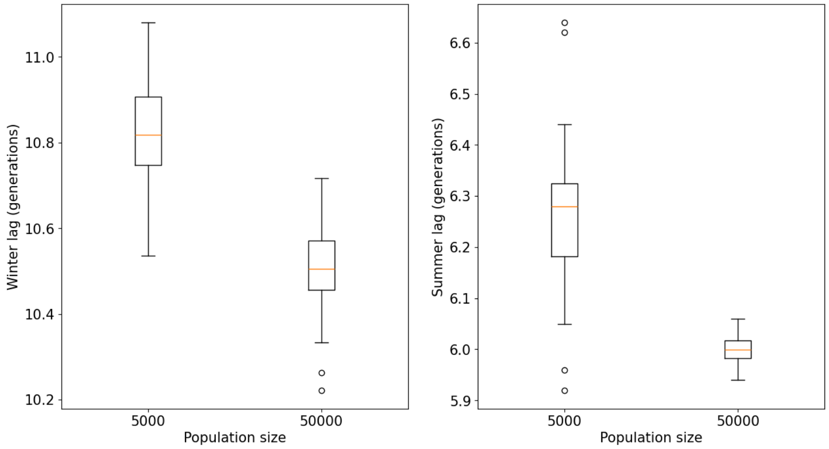


Figure S14: Distribution of winter and summer lags in the replicate simulations from Figure S13 (including seed=0 to 29). $N = 500$ was excluded since peaks and valleys are less easily identifiable.

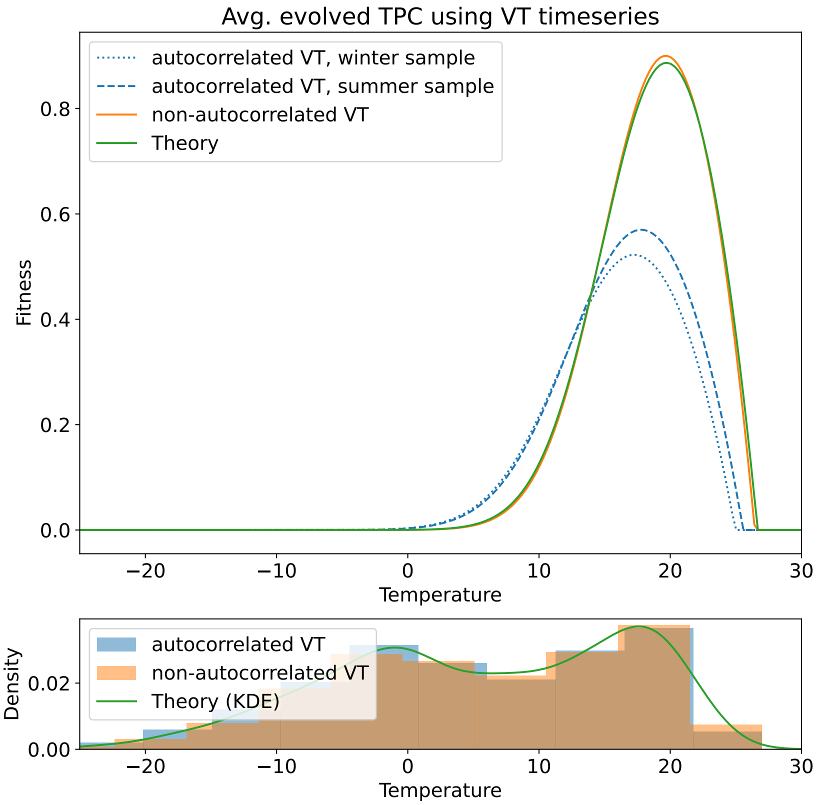


Figure S15: Simulated and theoretically expected TPC when temperature fluctuates according to NASA POWER data in Vermont (longitude=-72.47167, latitude=44.36272). First simulation (blue) uses the data as is, and average TPC was sampled from winter (blue dotted line in top panel) and summer (blue dash line in top panel) before the end of simulation. We plot temperature timeseries and the timing of two sampling events in Figure S16. Second simulation (orange) "scrambles" the timeseries data by sampling daily temperature from the timeseries data from the first simulation without order, and therefore removing autocorrelation. Finally, theoretically optimal TPC (green) was found by finding $B$ and $CTmin$ that maximizes $E[w_{\text{lifetime}}]$ using estimated probability density (KDE) from the VT timeseries data (green in the bottom panel). In other words, theory takes bimodal distribution into account (bottom panel; all three density distributions are approximately the same), and thus theoretical TPC matches simulated TPC when timeseries is randomized (green and orange in top). However, theory does not account for autocorrelation in temperature timeseries data, so simulated TPCs (blue in top panel; $B = 23.1$ in winter, $B = 23.3$ in summer) without scrambling are broader than theory (green in top panel; $B = 20.9$).

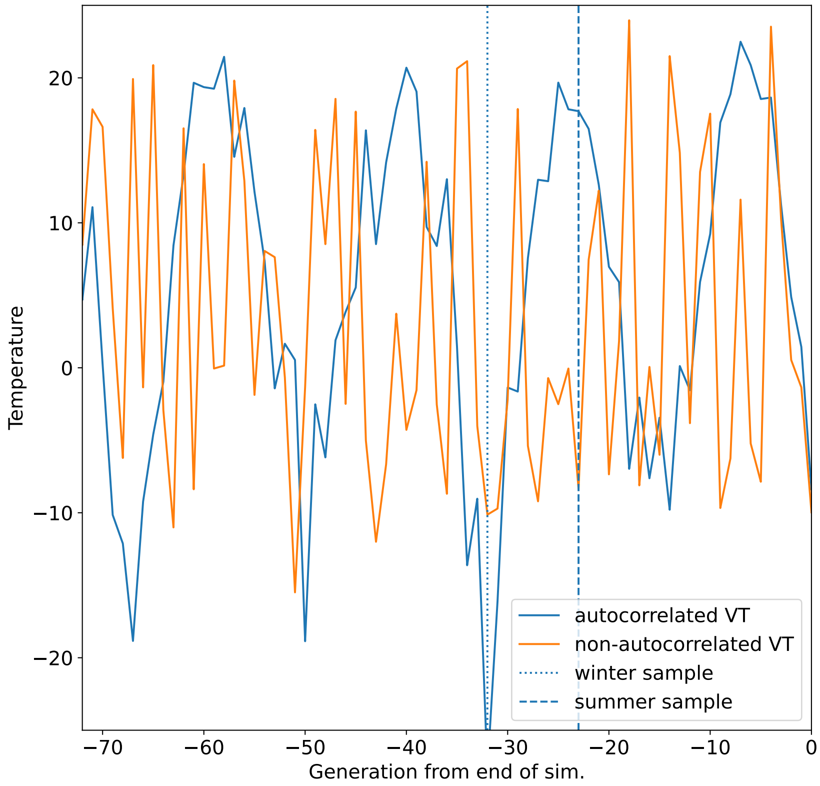


Figure S16: Temperatures (specifically, temperature on the first day of each generation) used in the simulation in the two cases plotted in Figure S15. Vertical lines mark when average TPC in Figure S15 is obtained. Original timeseries have high autocorrelation, while the scrambled data has minimal autocorrelation as every daily temperature is sampled independently from each other.

Table S1: Number of segregating QTNs for $B$ and $CTmin$ and their effect sizes ($\beta$) at the end of simulations that use Gaussian temperature distribution as in Figure 2.

| mean(T) | sd(T) | number of QTNs for $B$ | mean($\beta_B$) | std($\beta_B$) | number of QTNs for $CTmin$ | mean($\beta_{CTmin}$) | std($\beta_{CTmin}$) |
|---|---|---|---|---|---|---|---|
| 5 | 1 | 408 | -0.029892 | 0.208303 | 348 | -0.018062 | 0.181288 |
| 5 | 3 | 422 | -0.017649 | 0.218915 | 358 | -0.011388 | 0.188086 |
| 20 | 1 | 497 | -0.007703 | 0.215919 | 435 | -0.017340 | 0.194047 |
| 20 | 3 | 521 | -0.004768 | 0.203025 | 454 | -0.003597 | 0.185608 |
| 35 | 1 | 389 | -0.012341 | 0.189480 | 366 | 0.026760 | 0.183951 |
| 35 | 3 | 299 | -0.006246 | 0.191943 | 305 | 0.033317 | 0.175667 |

Table S2: Equilibration times ($t_{\text{equil}}$) measured for the simulations used in Figure S3 and Figure S4. We measure the time when the mean fitness has reached the 99 percent of the total increment between right after the burn-in period and the final generation.

| mean(T) | std(T) | $B_{\text{default}}$ | $CTmin_{\text{default}}$ | mean($t_{\text{equil}}$) | std($t_{\text{equil}}$) |
|---|---|---|---|---|---|
| 5 | 1 | 31 | 5 | 5880 | 181 |
| 5 | 3 | 31 | 5 | 5630 | 164 |
| 20 | 1 | 31 | 5 | 6044 | 214 |
| 20 | 3 | 31 | 5 | 5494 | 94 |
| 35 | 1 | 31 | 5 | 15908 | 1295 |
| 35 | 3 | 31 | 5 | 23593 | 3397 |
| 5 | 1 | 5 | 33 | 6130 | 289 |
| 5 | 3 | 5 | 33 | 8569 | 1429 |
| 20 | 1 | 5 | 33 | 6909 | 380 |
| 20 | 3 | 5 | 33 | 6574 | 302 |
| 35 | 1 | 5 | 33 | 5641 | 125 |
| 35 | 3 | 5 | 33 | 7218 | 1089 |

Table S3: Proportion of individuals with extremely high $CTmax$ ($r_{red}$), extremely low $CTmin$ ($r_{blue}$) individuals from the simulations in Figure 2, measured using all 30 replicate simulations for each treatment and using csv file of sample $B$ and $CTmin$ of 100 individuals in the final 100 generations. An individual is classified as "extremely high $CTmax$ if its $CTmax \geq CTmax_{\text{extreme}}$ (i.e. $w_{CTmax} < 0.5$) and as "extremely low $CTmin$" if $CTmin \leq CTmin_{\text{extreme}}$ (i.e. $w_{CTmin} < 0.5$). The individuals that have neither extreme $CTmax$ or $CTmin$ had none of the three fitness components for physiological costs ($w_B, w_{CTmin}, w_{CTmax}$) more extreme than the threshold (i.e. colored grey in Figure 2)

| | | $r_{blue}$ | | $r_{red}$ | |
|---|---|---|---|---|---|
| | | mean | std | mean | std |
| mean(T) | std(T) | | | | |
| 5 | 1 | 0.026330 | 0.002763 | 0.000000 | 0.000000 |
| | 3 | 0.027558 | 0.003396 | 0.000000 | 0.000000 |
| 20 | 1 | 0.000000 | 0.000000 | 0.000000 | 0.000000 |
| | 3 | 0.000000 | 0.000000 | 0.000000 | 0.000000 |
| 35 | 1 | 0.000000 | 0.000000 | 0.055224 | 0.003382 |
| | 3 | 0.000000 | 0.000000 | 0.129868 | 0.004674 |

[5] P. Virtanen, R. Gommers, T. E. Oliphant, M. Haberland, T. Reddy, D. Cournapeau, E. Burovski, P. Peterson, W. Weckesser, J. Bright, S. J. Van Der Walt, M. Brett, J. Wilson, K. J. Millman, N. Mayorov, A. R. J. Nelson, E. Jones, R. Kern, E. Larson, C. J. Carey, İ. Polat, Y. Feng, E. W. Moore, J. VanderPlas, D. Laxalde, J. Perktold, R. Cimrman, I. Henriksen, E. A. Quintero, C. R. Harris, A. M. Archibald, A. H. Ribeiro, F. Pedregosa, P. Van Mulbregt, SciPy 1.0 Contributors, A. Vijaykumar, A. P. Bardelli, A. Rothberg, A. Hilboll, A. Kloeckner, A. Scopatz, A. Lee, A. Rokem, C. N. Woods, C. Fulton, C. Masson, C. Häggström, C. Fitzgerald, D. A. Nicholson, D. R. Hagen, D. V. Pasechnik, E. Olivetti, E. Martin, E. Wieser, F. Silva, F. Lenders, F. Wilhelm, G. Young, G. A. Price, G.-L. Ingold, G. E. Allen, G. R. Lee, H. Audren, I. Probst, J. P. Dietrich, J. Silterra, J. T. Webber, J. Slavič, J. Nothman, J. Buchner, J. Kulick, J. L. Schönberger, J. V. De Miranda Cardoso, J. Reimer, J. Harrington, J. L. C. Rodríguez, J. Nunez-Iglesias, J. Kuczynski, K. Tritz, M. Thoma, M. Newville, M. Kümmerer, M. Bolingbroke, M. Tartre, M. Pak, N. J. Smith, N. Nowaczyk, N. Shebanov, O. Pavlyk, P. A. Brodtkorb, P. Lee, R. T. McGibbon, R. Feldbauer, S. Lewis, S. Tygier, S. Sievert, S. Vigna, S. Peterson, S. More, T. Pudlik, T. Oshima, T. J. Pingel, T. P. Robitaille, T. Spura, T. R. Jones, T. Cera, T. Leslie, T. Zito, T. Krauss,

U. Upadhyay, Y. O. Halchenko, and Y. Vázquez-Baeza. Scipy 1.0: fundamental algorithms for scientific computing in python. *Nature Methods*, 17:261–272, 2020. doi: 10.1038/s41592-019-0686-2.
